# Virulence gene profiles and phylogeny of Shiga toxin-positive *Escherichia coli* strains isolated from FDA regulated foods during 2010-2017

**DOI:** 10.1101/461327

**Authors:** Narjol Gonzalez-Escalona, Julie Ann Kase

## Abstract

Illnesses caused by Shiga toxin-producing *Escherichia* coli (STECs) can be life threatening, such as hemolytic uremic syndrome (HUS). The STECs most frequently identified by USDA’s Microbiological Data Program (MDP) carried toxin gene subtypes *stx1a* and/or *stx2a*. Here we describe the genome sequences of 331 STECs isolated from foods regulated by the FDA 2010-2017, determining their genomic identity, serotype, sequence type, virulence potential, and prevalence of antimicrobial resistance. Isolates were selected from the MDP archive, routine food testing by field labs (ORA), food testing by a contract company, and our laboratory (ORS). Only 276 (83%) were confirmed as STECs by *in silico* analysis. Foods from which STECs were recovered included cilantro (6%), spinach (25%), lettuce (11%), and flour (9%). Phylogenetic analysis using core genome MLST revealed these STEC genomes were highly variable, with some clustering associated with ST types and serotypes. We detected 95 different sequence types (ST); several ST were previously associated with HUS: ST21 and ST29 (O26:H11), ST11 (O157:H7), ST33 (O91:H14), ST17 (O103:H2), and ST16 (O111:H-). *in silico* virulome analyses showed ~ 51% of these strains were potentially pathogenic [besides *stx* gene they also carried *eae* (25%) or 26% *subA* (26%)]. Virulence gene prevalence was also determined: *stx*1 only (19%) -variants a and c; *stx*2 only (66%) – variants a, b, c, d, e, and g; and *stx*1/*sxt*2 (15%). Our data form a new WGS database that can be used to support food safety investigations and monitor the recurrence/emergence of *E. coli* in foods.

**Importance:** Shiga toxin-producing *Escherichia* coli (STECs) are associated with foodborne outbreaks worldwide; however, surveillance has not previously included genomic analyses for phylogenetics, prevalence, or potential virulence. We constructed the first genomic database of isolates from FDA-regulated foods to help monitor the emergence of new pathogenic STECs. Although only ~30 STECs were isolated per year, 50% of these carried markers associated with pathogenesis either a combination of *eae* plus *stx*, or *subA* plus *stx*. Moreover, those strains also carried virulence genes associated with severe illnesses. Here we showed that WGS enabled comparisons across isolates to establish phylogeny, help in identification of antibiotic resistance by monitoring the presence of antimicrobial resistance genes, and determined the presence of known virulence genes that have been linked with illnesses. Future food safety investigations will benefit from improved source tracking and risk assessments made possible by these analyses and new WGS database.

## Introduction

Shiga toxin-producing *Escherichia* coli (STECs) have the potential to cause infections, from mild to life threatening outcomes such as hemolytic uremic syndrome (HUS). STECs causing HUS, hemorrhagic colitis and bloody diarrhea are known as enterohemorrhagic *E. coli* (EHEC). Among the most common EHECs are O157:H7, O26, O121, O103, O111, and O145. O157:H7 strains are responsible for most foodborne outbreaks in the last two decades (1) while non-O157 serogroups, O26, O121, O103, O111, and O145 are the second most common cause of EHEC foodborne infections in the US (2,3) and worldwide (4-7). Each year in the US, O157:H7 cause an approximately 95,000 cases with 2,150 hospitalizations, with non-O157 STECs responsible for an estimated 170,000 cases (3). These serotypes carry Shiga toxin genes (*stx1* and/or *stx2*) and there are at least 130 EHEC serotypes that have been recovered from human patients. The US Department of Agriculture Food Safety and Inspection Services (USDA FSIS) in 2011, declared O26 and five other non-O157 serogroups, O45, O103, O111, O121, and O145 as adulterants in ground beef and non-intact beef products, and in mid-2012 began testing for these pathogens in both domestic and imported beef trimmings (8).

In order to cause illness, STEC strains need a set of genes that allow them to attach, colonize, and produce and secrete Shiga toxin protein (9-12). Genes described for attachment and colonization include *eae* (intimin), other proteins present in the locus of enterocyte effacement (LEE) locus, T3SS effectors, as well as biofilm production, and other virulence genes that are usually located in a plasmid (e.g. *ehxA*), referred as the virulence plasmid to differentiate it from other possible plasmids that can be carried by the same strain as well (9,11,13). Although the precise role of *ehxA* in STEC pathogenesis remains to be elucidated, several studies indicate an association of *ehxA* in clinical disease since 1) *ehxA* was found to be produced by many STEC associated with diarrheal disease and HUS (14-16), and 2) serum samples from HUS patients have been shown to react specifically to *ehxA* (17). Some STECs do not carry *eae*, however they possess other genes believed to compensated for the lack of *eae* (e.g. *subA, saa* or *sat*) (18,19).

STECs can be transmitted by various means with food remaining the predominant transmission route (1). Among the illnesses caused by STECs in FDA regulated food products (FRFDA), fresh produce has been implicated in several outbreaks, as well as some other atypical commodities, such as flour (20). Leafy greens and other agricultural food crops are particularly susceptible to contamination since they are grown in close contact with the ground where runoff from livestock areas, particularly cattle, contaminated irrigation water, manure used as fertilizer, and the intrusion of wildlife into growing fields can occur (21). Many of these same items are consumed raw and possibly with little cleaning. Some noteworthy *E. coli* outbreaks reported by the Center of Disease and Control (CDC) in the US in the last 10 years are: 2009 - beef (O157:H7) and prepackaged cookie dough (O157:H7); 2010 cheese (O157:H7), romaine lettuce (O145) and beef (O157:H7); 2011 - romaine lettuce (O157:H7), Lebanon bologna (O157:H7), and in-shell hazelnuts (O157:H7); 2012 - spinach and spring mix blend (O157:H7), unknown source (O145), and raw clover sprouts (O26); 2013 ready-to-eat salads (O157:H7), and frozen food products (O121); 2014 - raw clover sprouts (O121), and ground beef (O157:H7); 2015 - rotisserie chicken salad (O157:H7), and Mexican-style restaurant chain (O26); 2016 - flour (O121 and O26), and alfalfa sprouts (O157); 2017 - leafy greens (O157:H7), and soy nut butter (O157:H7); in this year, 2018, there has been an outbreak link to romaine lettuce caused by O157:H7 (https://www.cdc.gov/ecoli/outbreaks.html).

Beyond the noted outbreaks, there have been several reports on STECs found in FRFDA (22,23). The most comprehensive survey was the USDA Microbiological Data Program (MDP) that collected domestic and imported fresh fruit and vegetable samples from primarily terminal markets and wholesale distribution centers from 2001-2012 (https://www.ams.usda.gov/datasets/mdp/mdp-program-data-and-reports). This program tested approximately up to 15,000 samples annually, and tested for the presence of *Salmonella*, *E. coli O157:*H7, and other STECs. STEC were found most frequently from spinach samples (0.5%), and of the 132 STECs isolated, 9% were found to carry *eae*. The most prevalent Shiga toxin variants found were *stx*1a (22%) and/or *stx*2a (56%) (23). However, little other information about the genome content of those strains is publicly available.

Whole genome sequencing (WGS) technology is reshaping food safety and food-borne illness investigations (24). The use of WGS is becoming more useful as the cost of bacterial genome sequencing decreases every year. WGS cost per bacterial sequence is now comparable to PFGE. There are many attractive attributes with regards to the use of WGS for analyzing food samples including the potential to identify all pathogens present in that sample (25). Among other applications of the use of WGS are: it can help in identifying genes that allow for resistance/survival or virulence of certain bacterial strains (26-28), can help in establishing phylogenetic relationships among old strains of STECs isolated from either clinical cases or environmental samples (7,29,30), and can further help in Identifying matches between environmental and outbreak strains during outbreaks scenarios (26, 30-32). Furthermore, using WGS can help in identifying matches among bacterial strains isolated from environmental samples in production facilities and may help locate contamination sources (33). It can also be extremely helpful in establishing mechanism of evolution among pathogens (34). For example the 2011 outbreak in Germany linked to fenugreek seeds caused by an *E. coli* strain with a genomic backbone and virulence traits of entero-aggregative *E. coli* (EAEC) but had acquired a *stx* phage (*stx*2a gene variant) and caused a more aggressive disease with high HUS rate cases (35,36). This event highlighted the high plasticity of the *E. coli* genomes and it constitutes a warning of the possible arise of more of this new “hybrid pathotype” strains.

Therefore, we wanted to further characterize and catalog historical strains of STECs isolated from FRFDA by performing WGS analysis of every STEC strains isolated by the MDP and other FDA surveillance programs, as well as, some FDA historical isolates. This work establishes is the first genomic database of FRFDA isolates which in turn, will allow improved surveillance for both recurrence and the emergence of new strains that might be impacting our food supply. A total of 296 presumptive STECs were isolated during 2010-2017, and 35 additional STECs were historical isolates from our collection. The 331 presumptive STEC strains were analyzed for virulence genes [encompassing all *E. coli* virulent types - STEC, entero-pathogenic *E. coli* (EPEC), entero-toxigenic *E. coli* (ETEC), entero-invasive *E. coli* (EIEC), and EAEC], *in silico* MLST, and antibiotic resistance genes. Finally, their phylogenetic relationships and diversity were determined by whole genome phylogeny analysis using an allele-based whole genome multilocus sequence analysis (MLST) or core genome MLST analysis (cgMLST).

## Materials and Methods

### Bacterial strains and media

*E. coli* (n = 331) presumptive Shiga toxin-positive strains used in this study are listed in supplementary table 1 (Table S1). Each strain was assigned a CFSAN number for future tracking. The FRFDA strains were isolated by us (n = 196), FDA Office of Regulatory Affairs (ORA) laboratories (n = 74), and a contracting lab (n = 63) during 2012-2017 in the US.

**Table 1.** Frequency of STEC isolated by food commodity.

| Commodities | No.<br>strains | % |
| --- | --- | --- |
| Spinach | 88 | 31.88 |
| Flour | 59 | 21.38 |
| Lettuce | 35 | 12.68 |
| Cilantro (coriander) | 32 | 11.59 |
| Cheese | 9 | 3.26 |
| Leafy greens | 8 | 2.90 |
| Kale | 7 | 2.54 |
| Basil | 6 | 2.17 |
| Pepper | 6 | 2.17 |
| Alfalfa sprouts | 3 | 1.09 |
| Cantaloupe | 3 | 1.09 |
| Parsley | 3 | 1.09 |
| Tomatoes | 3 | 1.09 |
| Creamy soy Nut butter | 2 | 0.72 |
| pizza dough dry mix | 2 | 0.72 |
| sprouts | 2 | 0.72 |
| Almond | 1 | 0.36 |
| oats animal feed | 1 | 0.36 |
| Celery | 1 | 0.36 |
| Clover sprouts | 1 | 0.36 |
| cucumbers | 1 | 0.36 |
| animal feed | 1 | 0.36 |
| enviromental | 2 | 0.72 |
| **Total** | **276** | **16.30** |

The *stx*-negative strains were eliminated from this analysis.

### DNA preparation

Genomic DNA from each strain was isolated from overnight cultures using the DNeasy Blood and Tissue Kit (QIAGEN, Valencia, CA), following the manufacturer’s instructions. The resultant DNA extract was stored at −20°C until used as a template for whole genome sequencing. The concentration was determined using a Qubit double-stranded DNA HS assay kit and a Qubit 2.0 fluorometer (Thermo Fisher Scientific, Waltham, MA), according to manufacturer’s instructions.

### Whole genome sequencing, contig assembly and annotation

The genomes of the strains were sequenced, using an Illumina MiSeq sequencer (Illumina, San Diego, CA), with the 2×250 bp pair-end chemistry according to manufacturer’s instructions, at approximately 80X average coverage. The genome libraries were constructed using the Nextera XT DNA sample prep kit (Illumina). Genomic sequence contigs were *de novo* assembled using default settings within CLC Genomics Workbench v9.5.2 (QIAGEN) with a minimum contig size threshold of 500 bp in length.

### *in silico* serotyping

The serotype of each strain analyzed in this study was confirmed using the genes deposited in the Center for Genomic Epidemiology (http://www.genomicepidemiology.org) for *E. coli* as part of their web-based serotyping tool (SerotypeFinder 1.1 - https://cge.cbs.dtu.dk/services/SerotypeFinder) (37,37,38). We used Ridom for performing batch screening of the genomes analyzed. Briefly, all the genes were divided into O-type (*wzx* and *wzy*) and H-type (*fli*C) genes in FASTA format (ex. All *wzx* alleles were in a single FASTA file), and used as task template. For the virulence screening, a project was created using all three task templates and each whole genome sequence was screened for the presence of each gene type (O-type or H-type gene). Results were similar to SerotypeFinder, and as done for the virulence genes previously, the data was now in a database and new alleles (if found) could be added to the task templates.

### *in silico* MLST phylogenetic analysis

The initial analysis and identification of the strains were performed using an *in silico E. coli* MLST approach, based on the information available at the *E. coli* MLST website (http://mlst.warwick.ac.uk/mlst/dbs/Ecoli) and using Ridom SeqSphere+ software v2.4.0 (Ridom; Münster, Germany) (http://www.ridom.com/seqsphere). Seven housekeeping genes (*dnaE, gyrB, recA, dtdS, pntA, pyrC,* and *tnaA*), described previously for *E. coli* (39), were used for MLST analysis. The same *E. coli* MLST database was also used to assign numbers for alleles and STs.

### *in silico* determination of virulence genes

Virulence genes were determined using the genes deposited in the Center for Genomic Epidemiology (http://www.genomicepidemiology.org) for *E. coli* as part of their VirulenceFinder 1.5 web-based tool (https://cge.cbs.dtu.dk/services/VirulenceFinder) (38), except that we used Ridom for performing batch screening of the genomes analyzed. Briefly, all the genes were divided into classes or groups by homology in FASTA format (e.g. All *ast*A alleles were in a single FASTA file), and used as a task template. Afterwards a project was created using all these task templates, and each WGS was screened for the presence of each gene class (virulence gene). We tested for 95 virulence genes previously reported here (27). These 95 virulence genes include different *E. coli* pathotypes (ETEC, STEC, EAEC, and EPEC) in order to detect any possible E. coli hybrid if present as observed for the O104:H4 Germany. The *stx* gene variants analyzed are available at https://cge.cbs.dtu.dk/services/VirulenceFinder. The result was very similar to the one displayed at VirulenceFinder, except that the data are now in a database and new alleles (if found) could be added to the task templates.

### *in silico* antimicrobial resistance genes identification

Antimicrobial resistance genes present in sequenced genomes as well as in those retrieved from GenBank (Table 2) were identified by using the genes deposited in the Center for Genomic Epidemiology (http://www.genomicepidemiology.org) as part of their Resfinder 2.1 web-based tool (https://cge.cbs.dtu.dk/services/ResFinder) (40), except that we used Ridom for performing batch screening of the genomes analyzed. Briefly, all the genes were divided into classes or groups by homology in fasta format (e.g. All *bla*TM alleles were located in a single fasta file), and used as task template. Later a project was created using all these task templates, and each WGS was screened for the presence of each gene class (antimicrobial resistance gene). The result was very similar to the one displayed at ResFinder, except that the data are now in a database and new alleles (if found) could be added to the task templates.

**Table 2.**
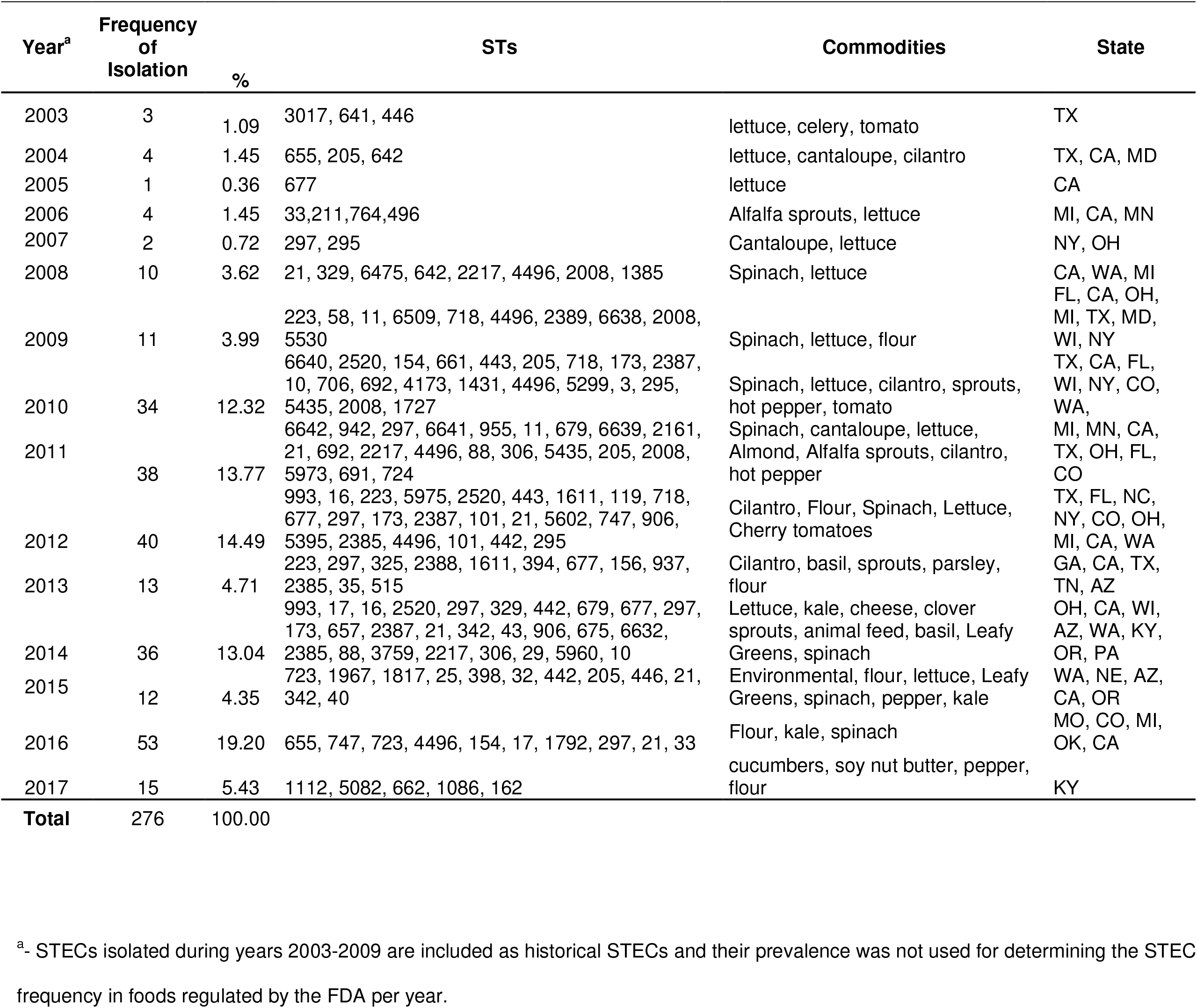
Frequency of STECs isolated from food commodities per year.

### Phylogenetic relationship of the strains by cgMLST analysis

The phylogenetic relationship of the strains was assessed by a core genome multilocus sequence typing (cgMLST) analysis using Ridom SeqSphere+ software v2.4.0. The genome of O26:H11 strain 11368 (NC_013361.1) was used as a reference. After eliminating loci that were missing from the genome of any strain used in our analyses, we performed a cgMLST analysis. These remaining loci were considered the core genome shared by the analyzed strains. We used Nei’s DNA distance method (41) for calculating the matrix of genetic distance, taking only the number of same/different alleles in the core genes into consideration. A Neighbor-Joining (NJ) tree using the appropriate genetic distances was built after the cgMLST analysis. The discriminatory index was calculated with the Ridom software using the Simpson’s discriminatory index as described (42); cgMLST uses alleles number of each loci for determining the genetic distance and build the phylogenetic tree. The use of allele numbers reduces the influence of recombination in the dataset studied and allow for fast clustering determination of genomes

### Nucleotide sequence accession numbers

The draft genome sequences of 196 *E. coli* strains used in our study are available in GenBank under the accession numbers listed in Table S1.

## RESULTS

### Presence of STEC in FDA regulated foods

Among 331 suspected STEC strains isolated from FRFDA between 2003 – 2017 and sequenced by several labs and deposited at NCBI, only 276 were confirmed to be STECs by *in silico* analysis for the presence of either *stx1* or *stx2* (Table S1). Of the 196 identified and sequenced by our lab, 92% carried either stx1 or 2 (181/196). From the 74 strains which genomes were retrieved from NCBI and were initially isolated and sequenced by FDA ORA, 94% carried either stx1 or 2 (70/74). Of the 63 *E. coli* strains isolated and sequenced by a FDA contracting laboratory, 43% carried either stx1 or 2 (25/61). The frequency of isolation of STECs from foods are listed in Table 1. STECs were isolated from 22 food commodities. Most of these STECs were isolated from spinach (32%), flour (21%), lettuce (13%), and cilantro (12%).

The frequency of STEC isolation per year, their sequence type, food commodity and state of isolation (if available) is listed in Table 2. A median of 30 STECs were recovered from FDA regulated foods per year. We used 2010 as our starting year, since the number of strains before that year were sporadically found and came from our STEC historical collection. The STEC strains analyzed were isolated in 22 states.

### Characterization of STEC strains by *in silico* MLST

Among the 276 STECs analyzed in this study we identified 95 different sequence types (STs) (35%) by *in silico* MLST. Strains belonging to a ST were isolated between 1 to 12 times in the period studied (2010-2017) (Table 3 and Table S2). The majority of the STs identified were observed only one time (45%).

**Table 3.**
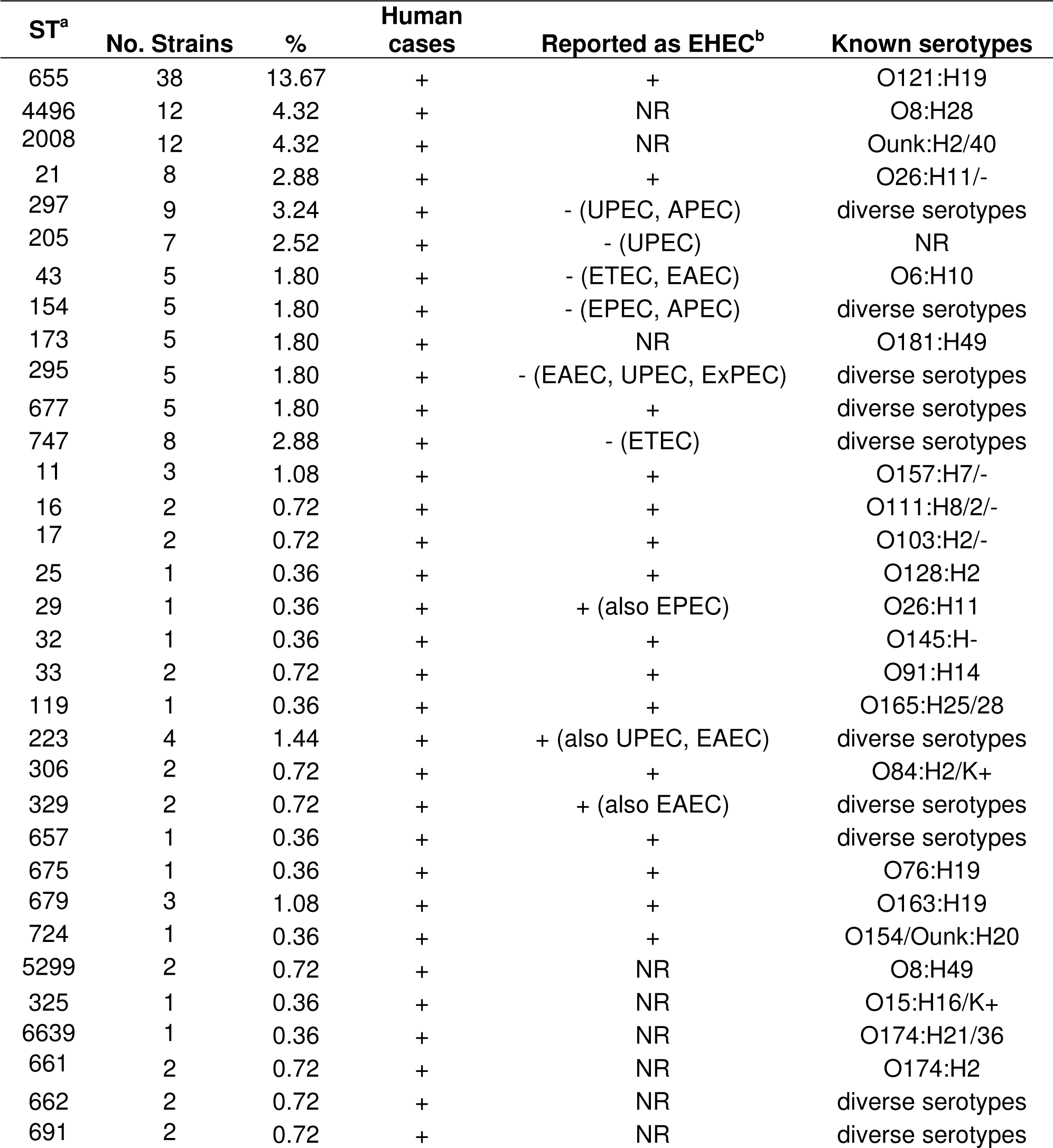
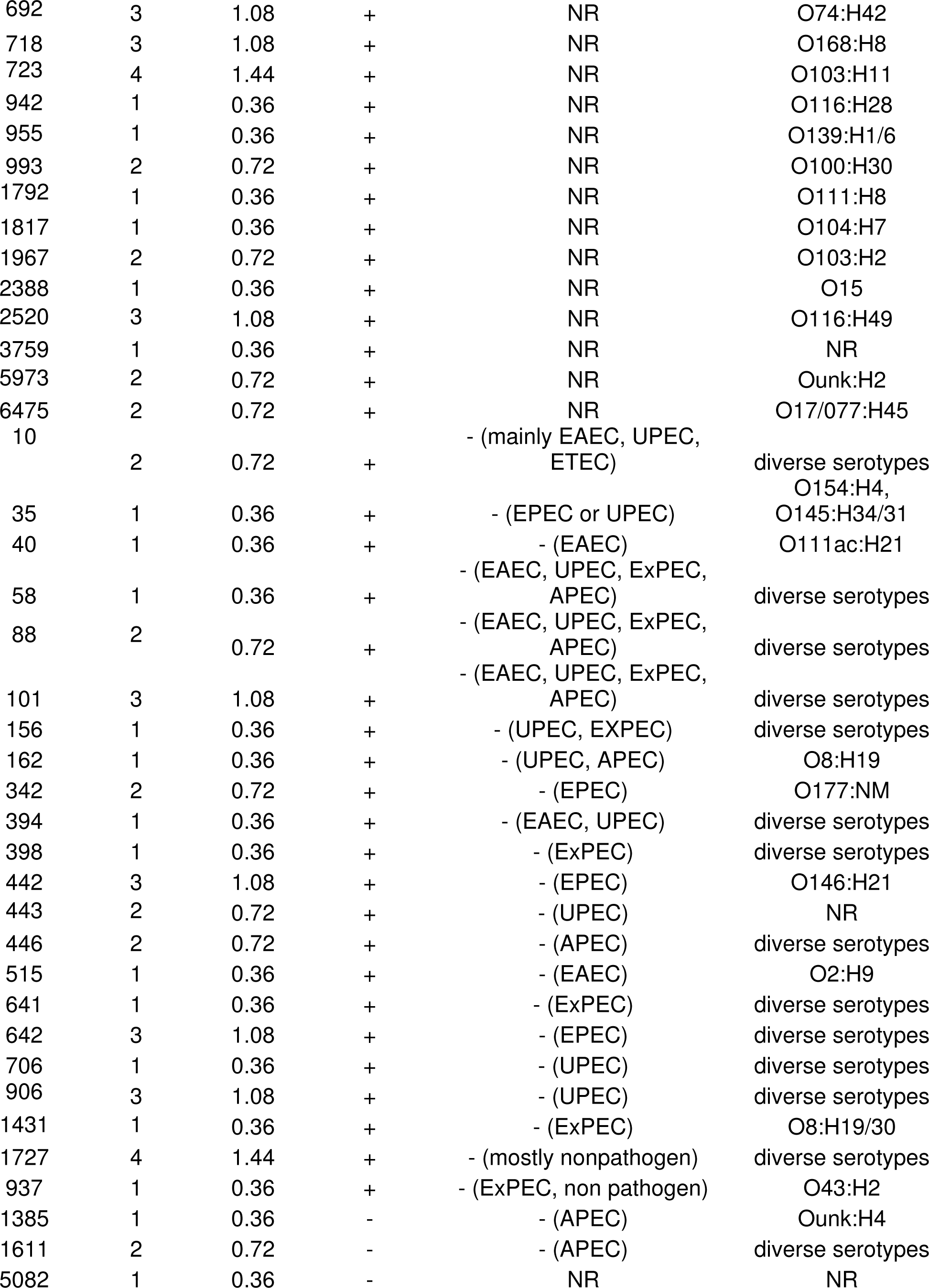
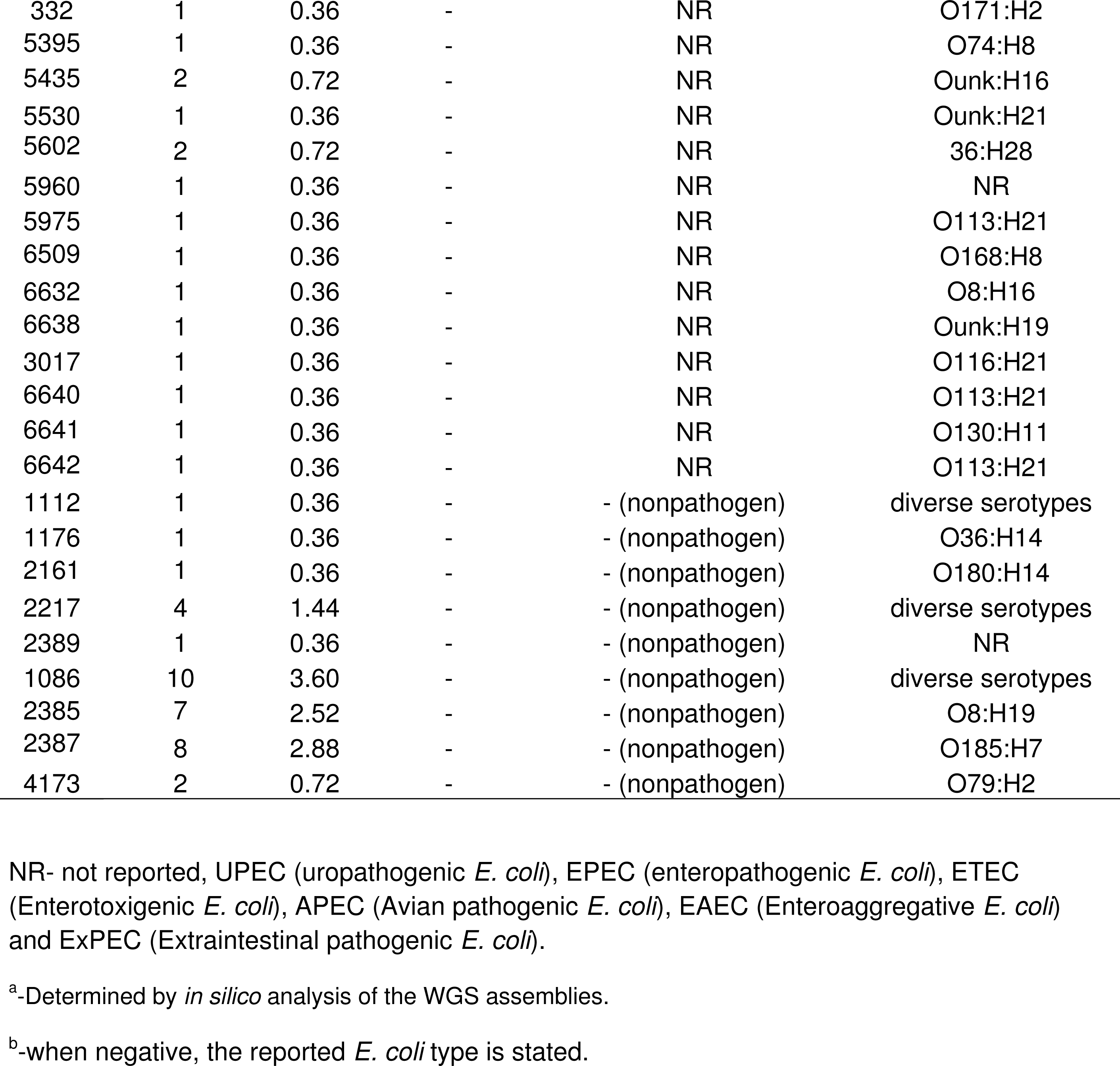
STs observed and number of strains included in each ST. Additionally information is provided for strains belonging to those STs such as: link to human cases, link to EHEC cases, and known serotypes. These additional reports are based on what it is reported in the *E. coli* section of the Enterobase database (http://enterobase.warwick.ac.uk).

### Characterization of STEC strains by serotyping, and virulence gene profiles

However, belonging to a known ST that caused HC is not enough to predict the probability of the strain to cause disease illness. Therefore, we further characterized these STECs by in silico virulence determination as well as their predicted serotype (Table 4). The detailed in silico analysis for presence of virulence genes and serotype is listed in Supplementary Table 2. Table 4 lists only the serotype and some of the most known virulence genes for each strain: *stx1* type*, stx2* type*, eae* type*, ehxA, espP, etpD, toxB, katP, subA, saa*, and *sab*. We identified at least 81 different serotypes among the 276 STECs sequenced (Table 4). Many of the O types were not present in our O types database and were listed as unknown. Among those some of the most common clinical STECs serotypes were identified, such as: O157:H7, O26:H11, O113:H21, O121:H19, O91:H21, O103:H2, and O111:H8.

**Table 4.**
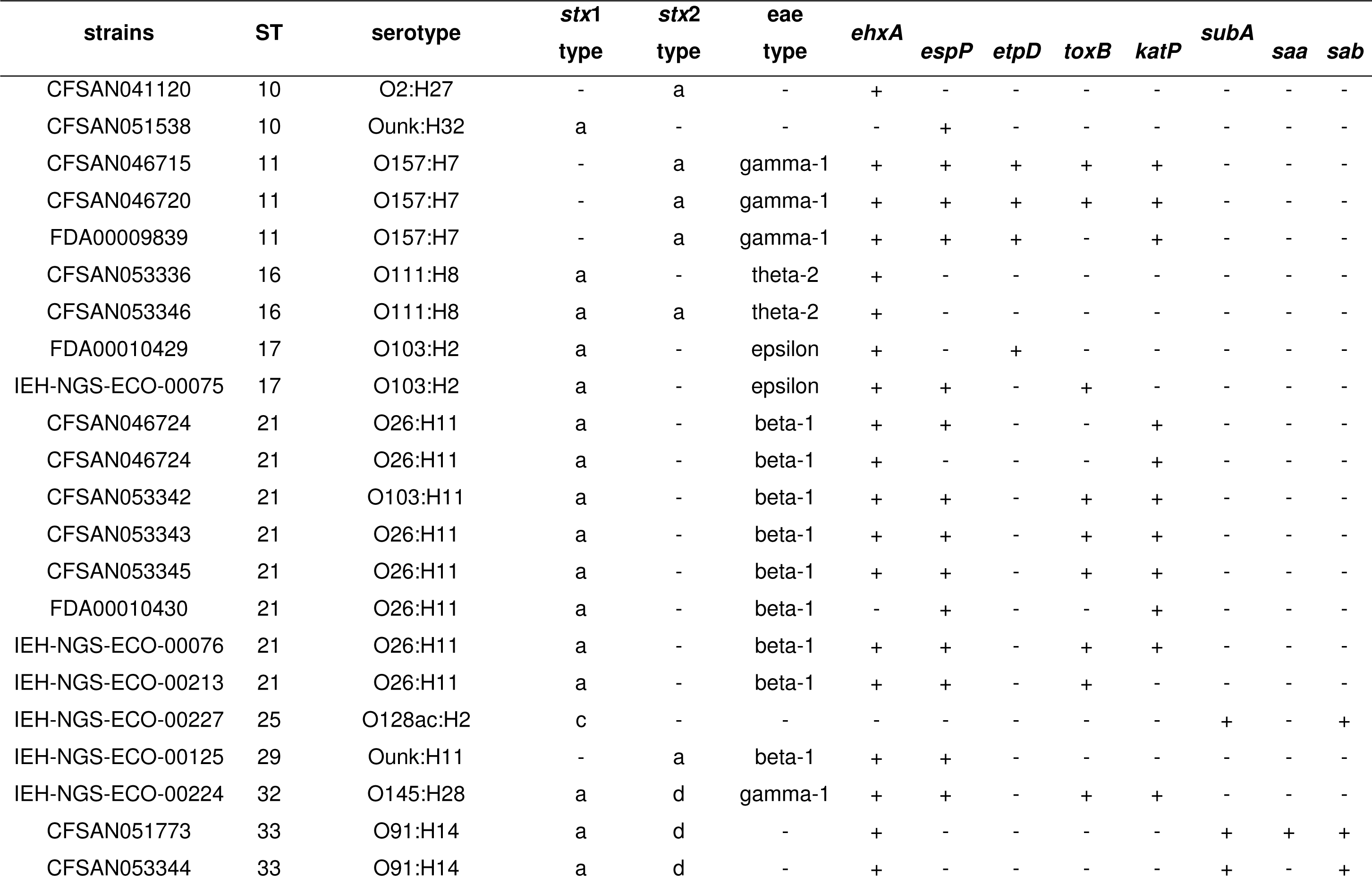
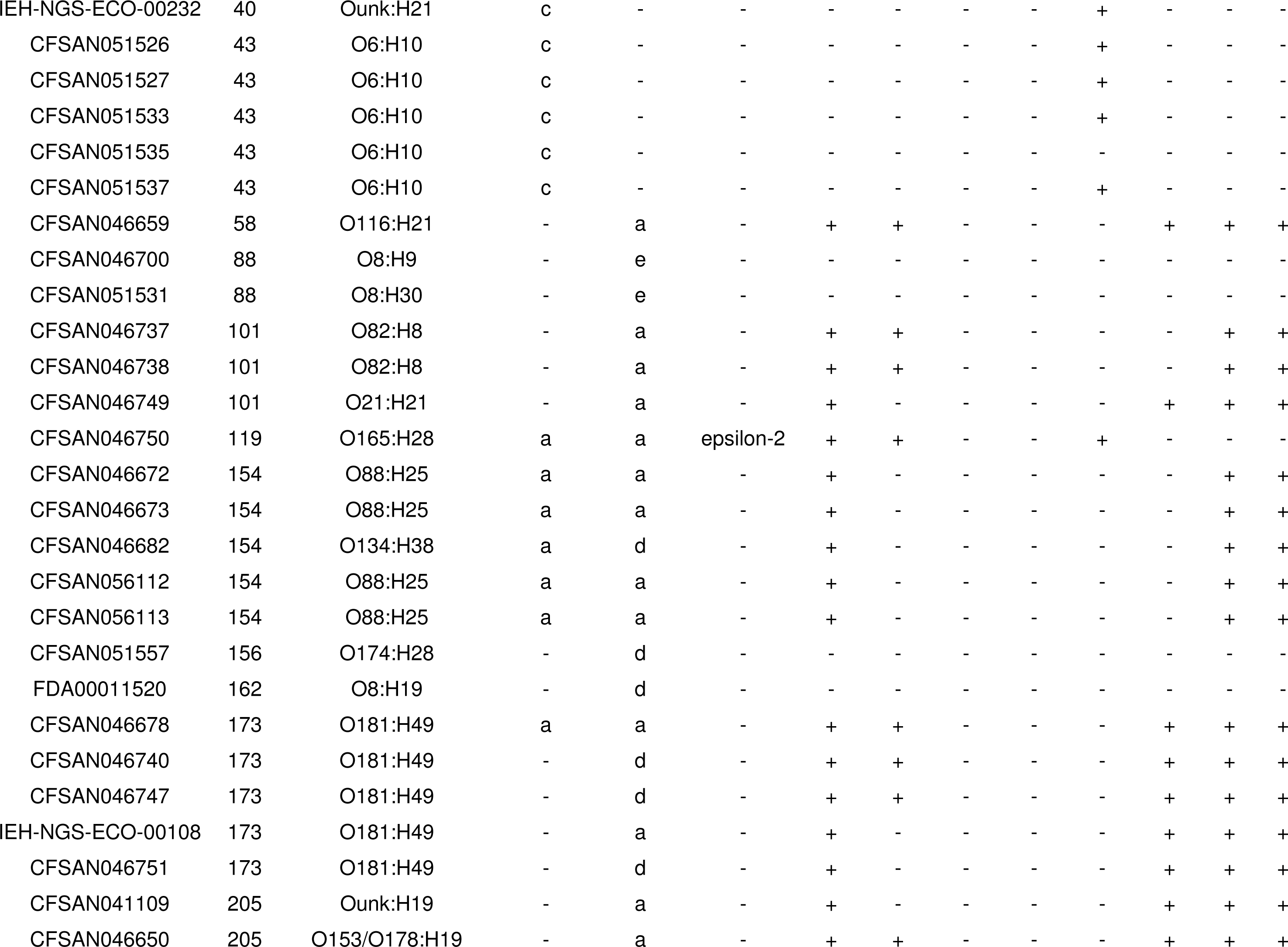
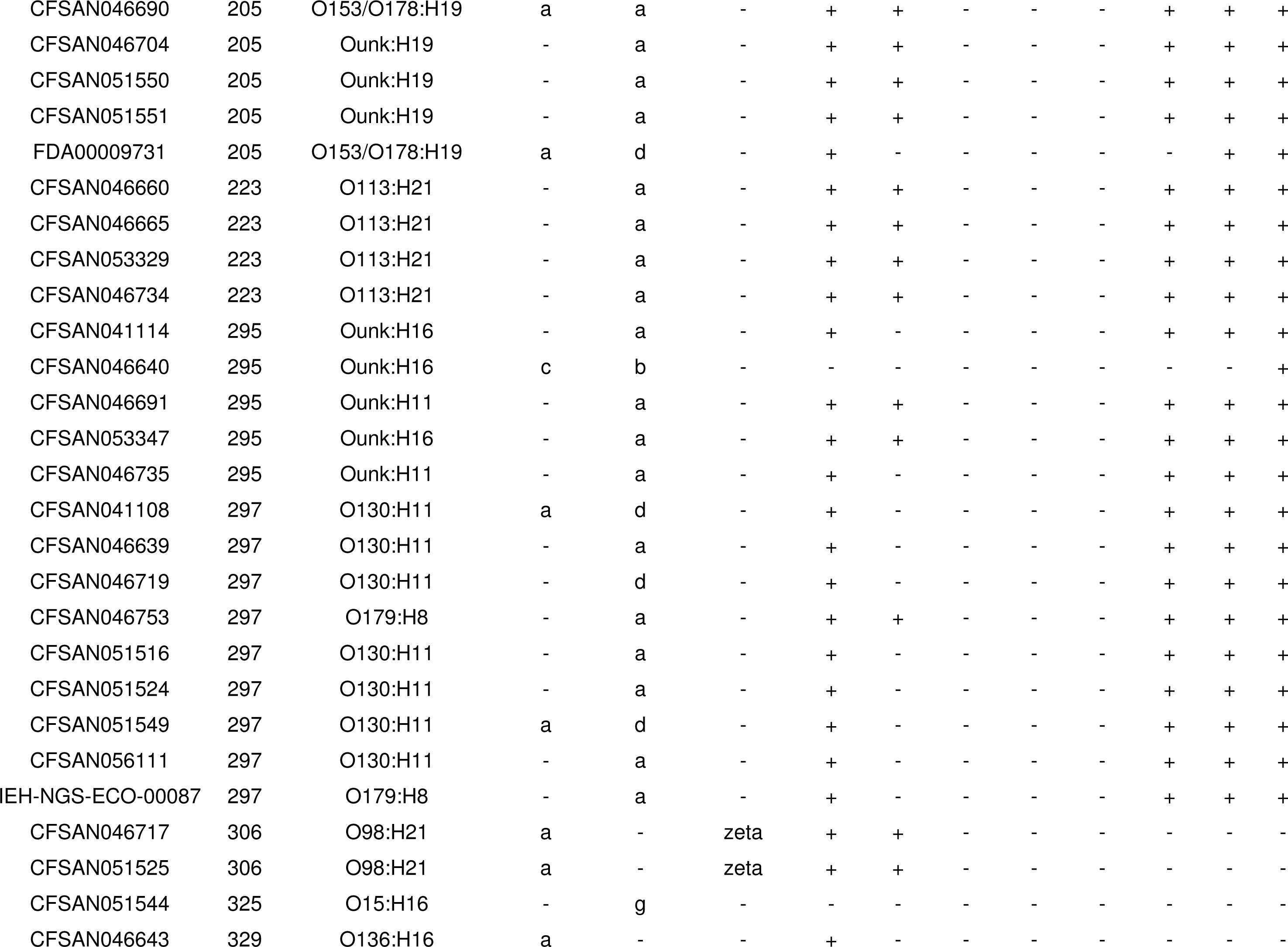
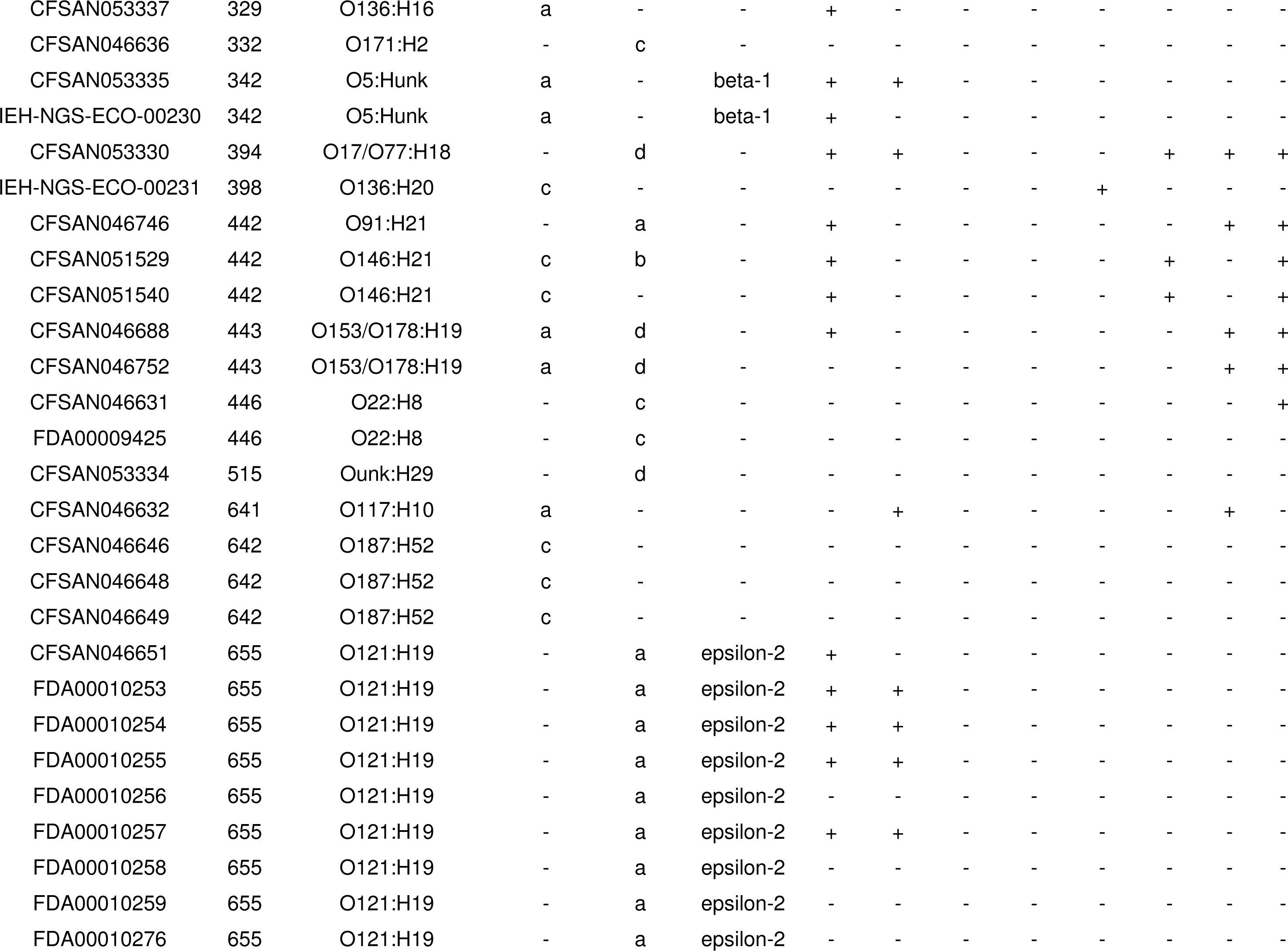
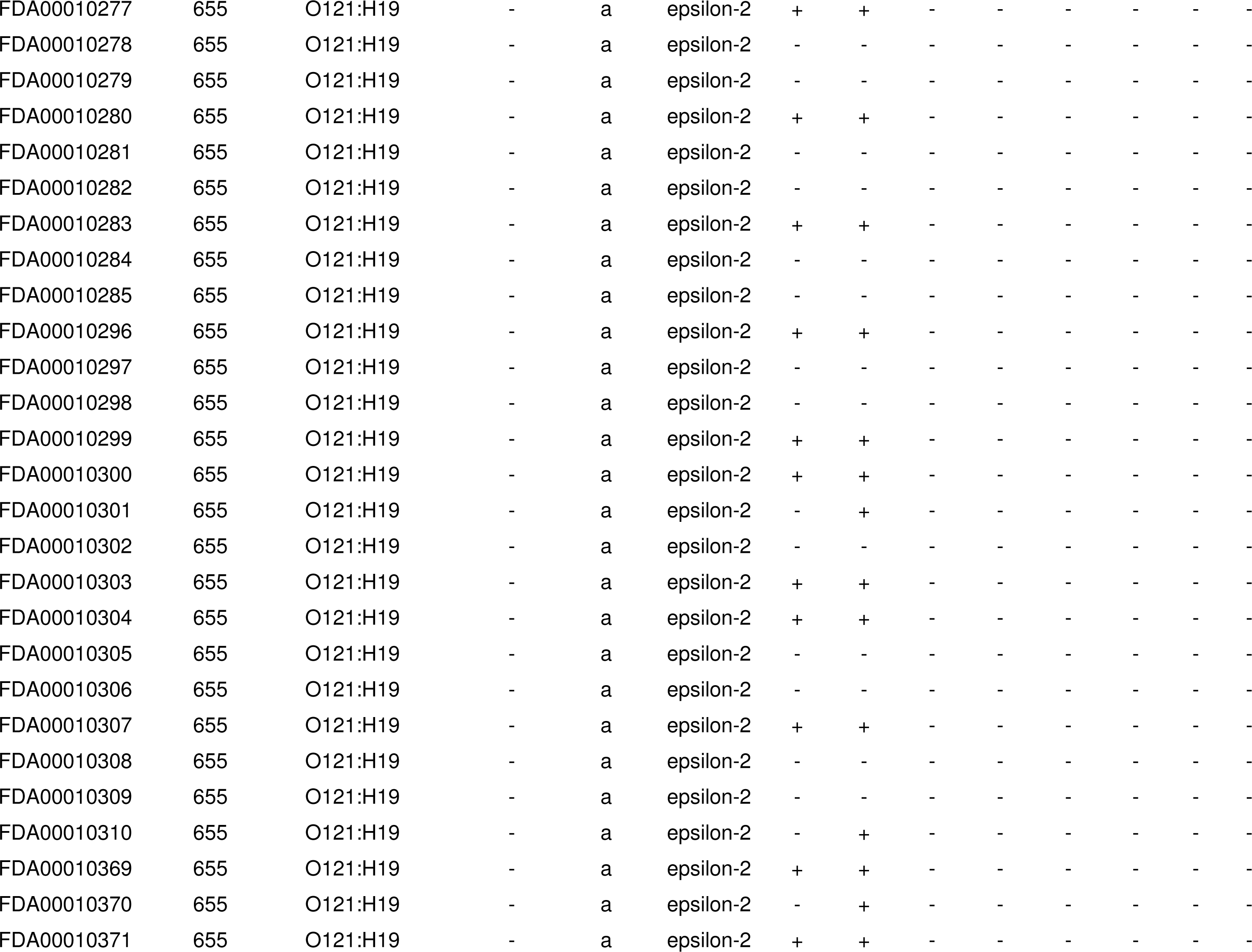
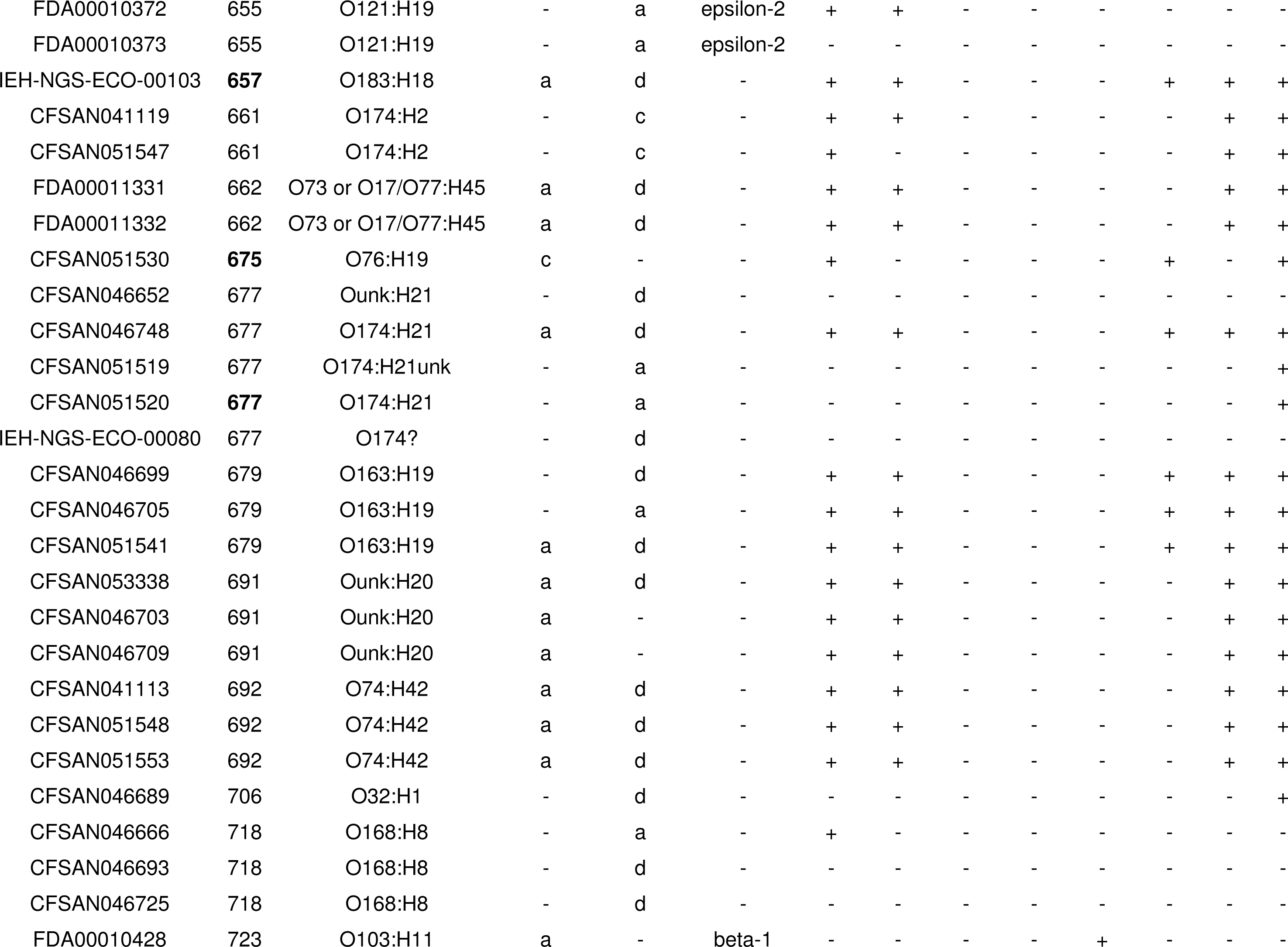
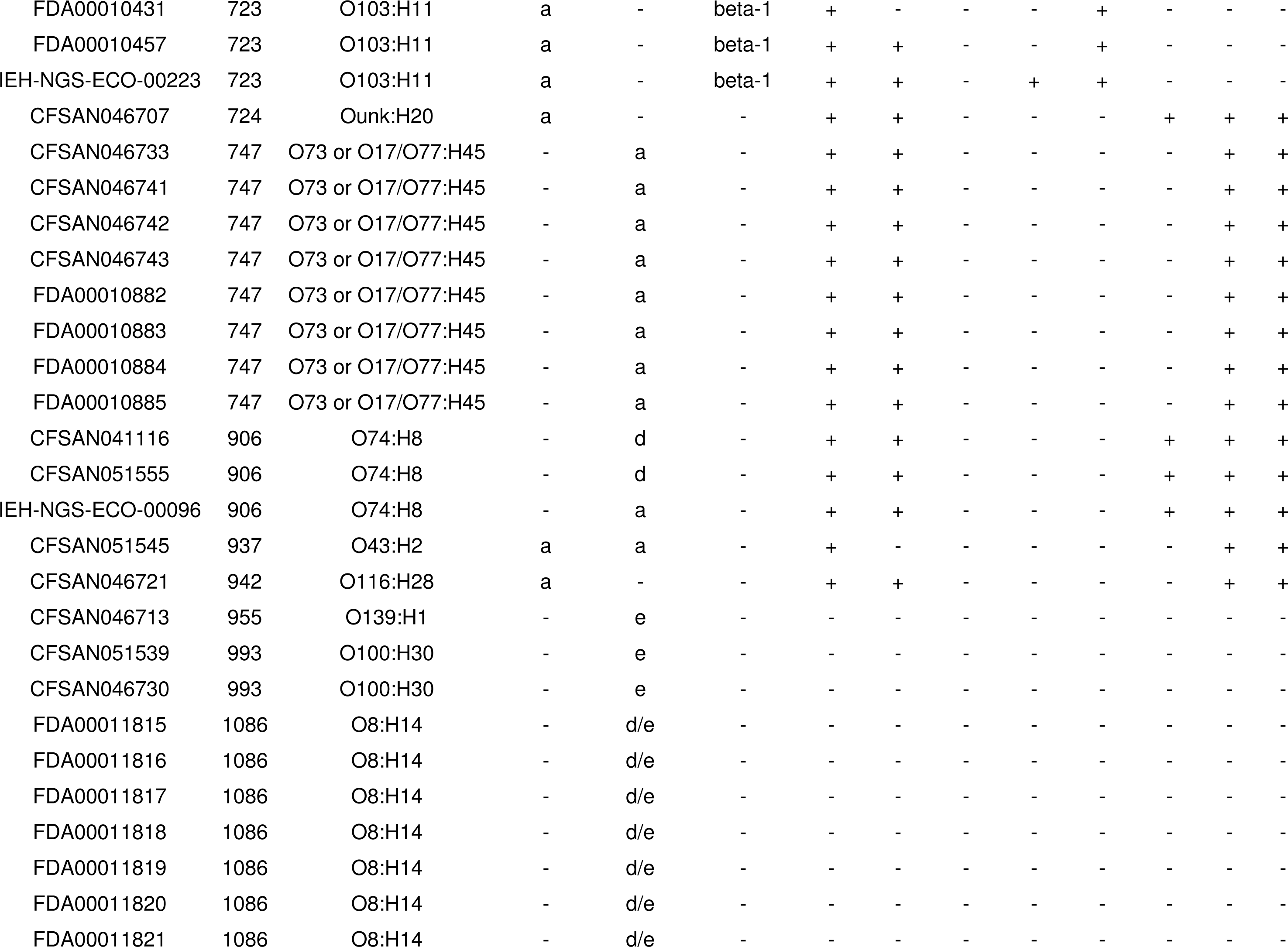
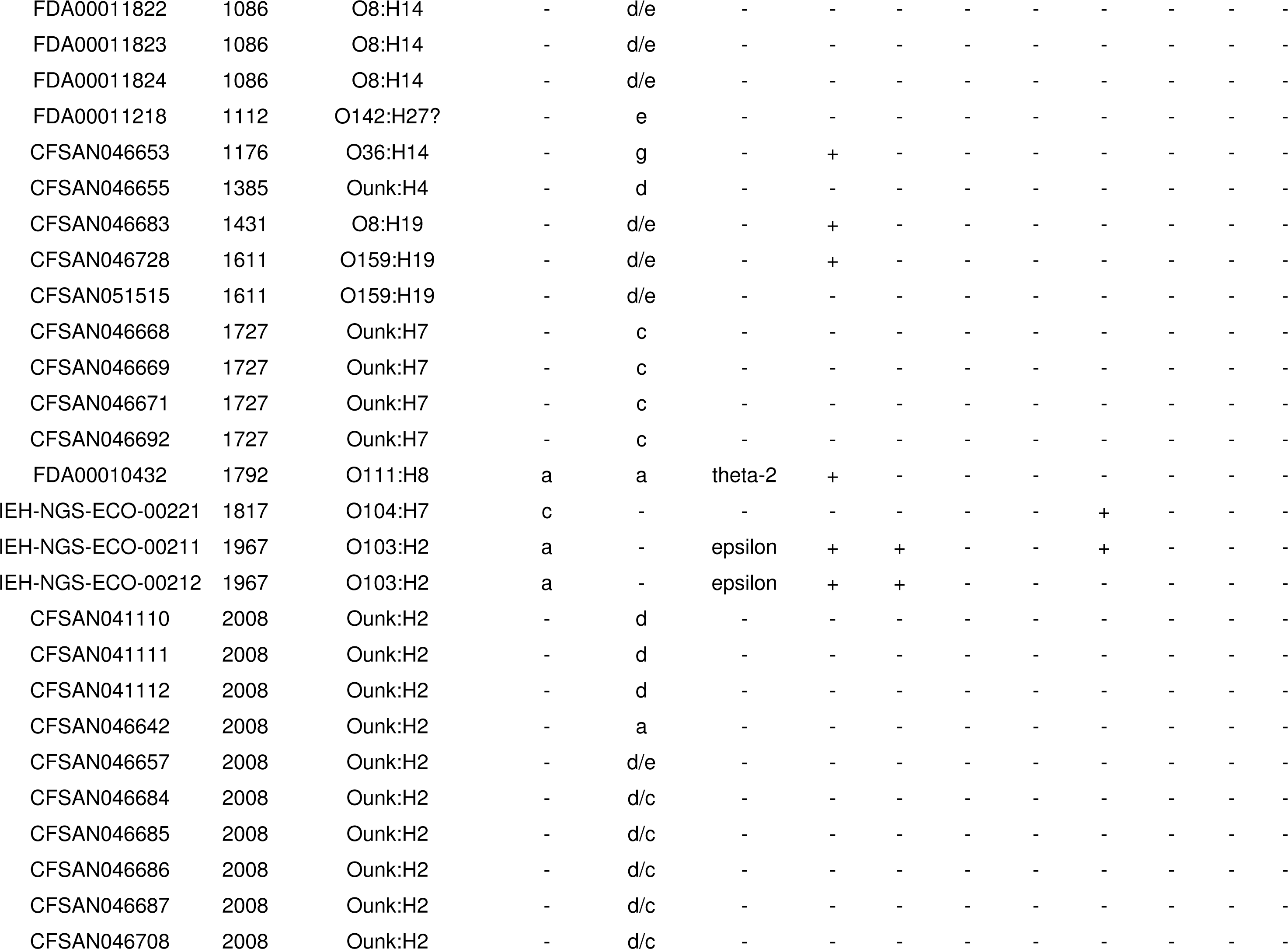
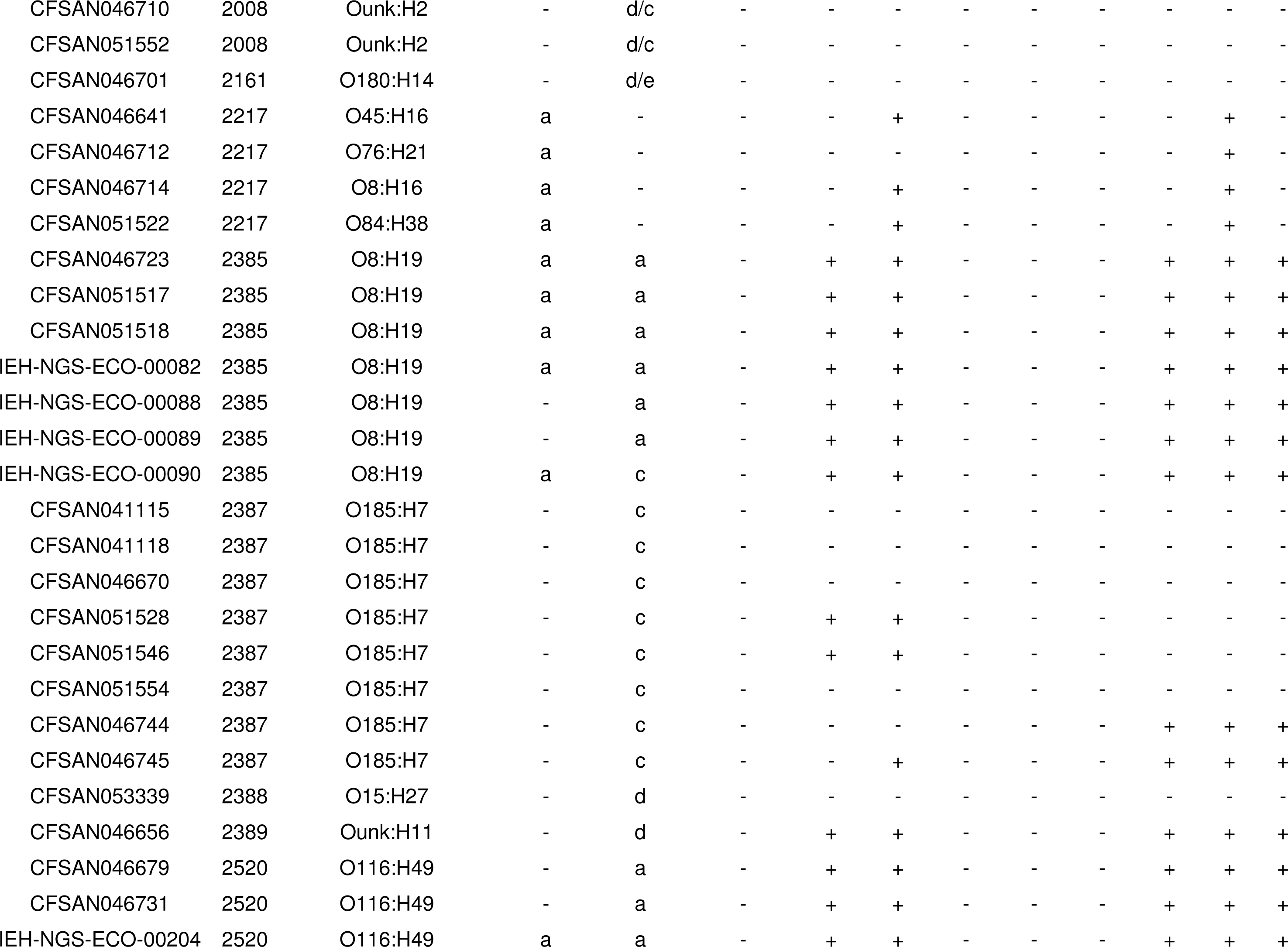
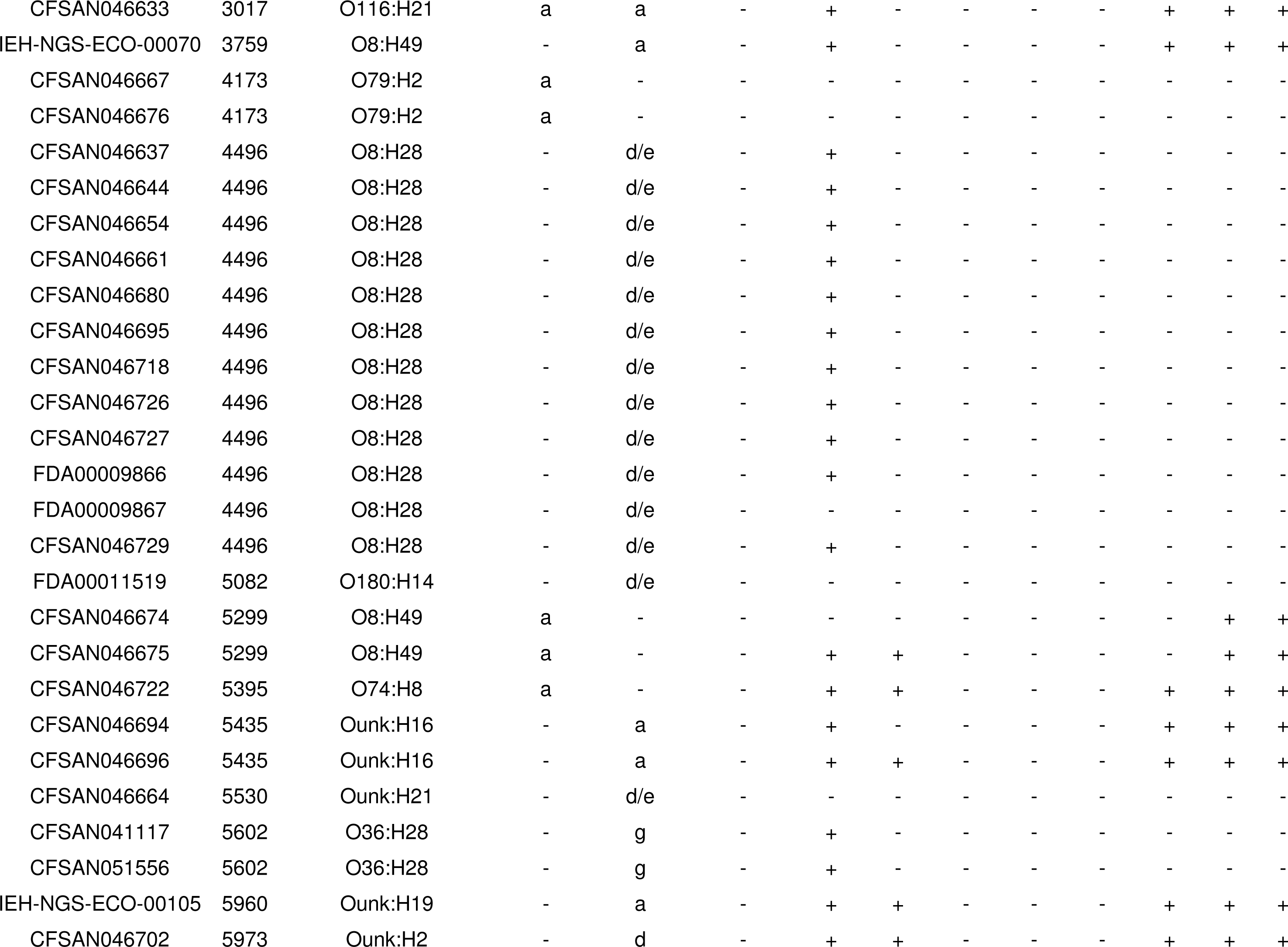
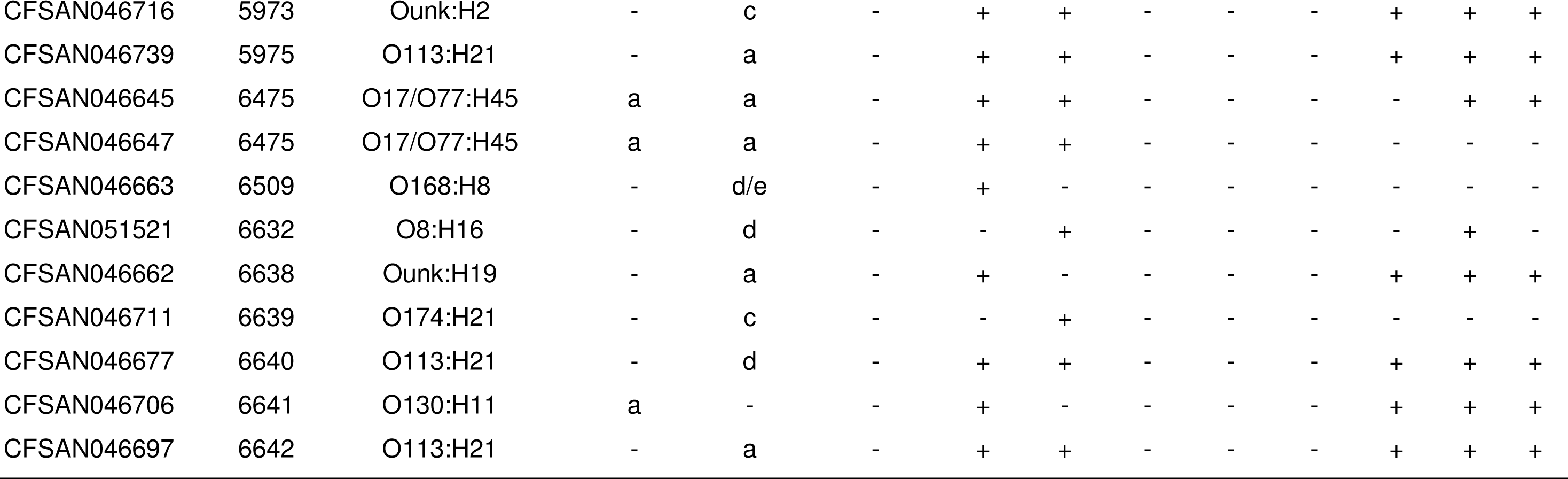
*in silico* characterization of STECs from this study for presence of virulence genes and their serotype.

Adherence factors *eae* and *subA* genes were found in 67 (24%) and 72 (26%), respectively (Table 4). Shiga toxin genes were present as follows: *stx*1- 53 (19%) (variants a and c), *stx*2- 184 (67%) (variants a, b, c, d, d/e, e, and g), while *stx*1+*stx*2 – 39 (15%). The other virulence genes were more sporadically found: *exhA* gene was present in 169 (61%), *espP* was present in 118 (43%), *katP* in 24 (9%), *etpD* in 4 (2%), and finally *toxB* was present in 10 (4%).

### Presence of antimicrobial resistance genes

Thirty-three of the 276 STEC strains (12%) carried antimicrobial resistance genes (Table 5). Thirty of them carried multiple antibiotic resistance genes while the remainder three carried a single gene (*tetA*- IEH-NGS-ECO-00231, FDA00011218, and CFSAN051521). Among the antimicrobial classes observed were genes resistant to aminoglycosides, beta-lactamases, macrolides, phenicols, quinolones, sulphonamides, tetracyclines, and trimethoprim.

**Table 5.** Presence of antimicrobial resistance genes identified by *in silico* analysis in the 276 STEC genomes analyzed in this study.

| Strains | *aadA*<sup>a</sup> | *aph3*<sup>a</sup> | *strA*<sup>a</sup> | *strB*<sup>a</sup> | *bla*<br>*TEM*<sup>b</sup> | *floR*<sup>d</sup> | *QnrB*<sup>e</sup> | *sul1*<sup>f</sup> | *sul2*<sup>f</sup> | *sul3*<sup>f</sup> | *tetA*<sup>g</sup> | *tetB*<sup>g</sup> | *tetC*<sup>g</sup> | *dfrA*<sup>ha</sup> |
| --- | --- | --- | --- | --- | --- | --- | --- | --- | --- | --- | --- | --- | --- | --- |
| CFSAN053338 | - | - | + | + | - | - | - | - | - | - | - | - | - | - |
| CFSAN046714<sup>g</sup> | - | - | - | - | - | - | - | + | - | + | - | - | + | - |
| CFSAN051552 | - | - | + | + | - | - | - | - | + | - | - | + | - | - |
| CFSAN046710 | - | - | + | + | - | - | - | - | + | - | - | + | - | - |
| CFSAN046642 | - | - | + | + | - | - | - | - | + | - | - | + | - | - |
| IEH-NGS-ECO-00231 | - | - | - | - | - | - | - | - | - | - | + | - | - | - |
| CFSAN053336 | - | + | + | + | - | - | - | - | + | - | + | - | - | - |
| FDA00009425 | - | - | + | + | - | + | - | - | + | - | + | - | - | - |
| FDA00011218 | - | - | - | - | - | - | - | - | - | - | + | - | - | - |
| IEH-NGS-ECO-00230<sup>i</sup> | + | - | + | + | - | + | - | + | + | + | - | - | - | - |
| CFSAN051526 | + | - | - | - | - | - | - | + | - | + | + | - | - | - |
| CFSAN046636 | - | - | + | + | - | - | - | - | + | - | - | + | - | - |
| CFSAN046669 | - | - | + | + | - | - | - | - | + | - | - | + | - | - |
| CFSAN046668 | - | - | + | + | - | - | - | - | + | - | - | + | - | - |
| CFSAN053334 | - | - | + | + | - | + | - | - | + | - | + | - | - | - |
| CFSAN046687 | - | - | + | + | - | - | - | - | + | - | - | + | - | - |
| CFSAN051521 | - | - | - | - | - | - | - | - | - | - | + | - | - | - |
| CFSAN046730 | - | + | + | + | - | - | - | - | + | - | - | + | - | - |
| CFSAN041112 | - | - | + | + | - | - | - | - | + | - | - | + | - | - |
| CFSAN041111 | - | - | + | + | - | - | - | - | + | - | - | + | - | - |
| CFSAN046671 | - | - | + | + | - | - | - | - | + | - | - | + | - | - |
| CFSAN046708 | - | - | + | + | - | - | - | - | + | - | - | + | - | - |
| CFSAN046685 | - | - | + | + | - | - | - | - | + | - | - | + | - | - |
| CFSAN046684 | - | - | + | + | - | - | - | - | + | - | - | + | - | - |
| CFSAN051527 | + | - | - | - | - | - | - | + | - | + | + | - | - | - |
| CFSAN046686 | - | - | + | + | - | - | - | - | + | - | - | + | - | - |
| CFSAN051535 | - | - | + | + | + | - | - | - | + | - | + | - | - | - |
| CFSAN046693 | - | - | + | + | - | - | - | - | + | - | - | - | - | - |
| CFSAN046713 | - | - | - | + | + | - | + | - | + | - | + | - | - | + |
| CFSAN046725 | - | - | - | + | - | - | - | - | + | - | - | - | - | - |
| CFSAN051531 | - | - | - | + | - | - | - | - | + | - | + | - | - | + |
| CFSAN051539<sup>j</sup> | + | + | - | - | + | - | - | - | - | + | - | + | - | - |
| CFSAN041110 | - | - | + | + | - | - | - | - | + | - | - | + | - | - |

<sup>a</sup>Aminoglycoside, <sup>b</sup>Beta-lactamase, <sup>c</sup>Macrolide, <sup>d</sup>Phenicol, <sup>e</sup>Quinolone, <sup>f</sup>Sulphonamide, <sup>g</sup>Tetracycline, and <sup>h</sup>Trimethoprim.
<sup>g</sup> strain carrying *blaOXA*<sup>b</sup> gene. <sup>i</sup> strain carrying *blaCMY-2*<sup>b</sup> gene. <sup>j</sup> strain carrying *mef(B)*<sup>c</sup>, *cmf*<sup>d</sup> and *cmIA1*<sup>d</sup> genes.

### Phylogenetic relationship of the STEC strains by cgMLST analysis

The phylogenetic relationships among the 276 STECs from this study determined by cgMLST analysis is shown in Figure 1. The genome of O157:H7 strain Sakai (NC_002695.1) was used as the reference for the cgMLST. This *E. coli* strain has 5,204 genes, of which 3,860 genes (core genes) were present in the six genomes used as comparison to generate the cgMLST scheme (NC_011353.1 – O157:H7 strain EC4115, NC_002655.2 - O157:H7 strain EDL933, NC_013008.1 – O157:H7 strain TW14359, NC_013941.1 - O55:H7 strain CB9615, NC_017656.1 - O55:H7 strain RM12579, and NC_017906.1 – O157:H7 strain Xuzhou21). While 791 genes were found in some of the compared genomes. The remainder of the genes were eliminated from the analysis for several reasons (genes were paralogous, or pseudogenes). Therefore, a total of 4,651 genes were used as templates for the analysis of the STECs from this study.

**Figure 1.**
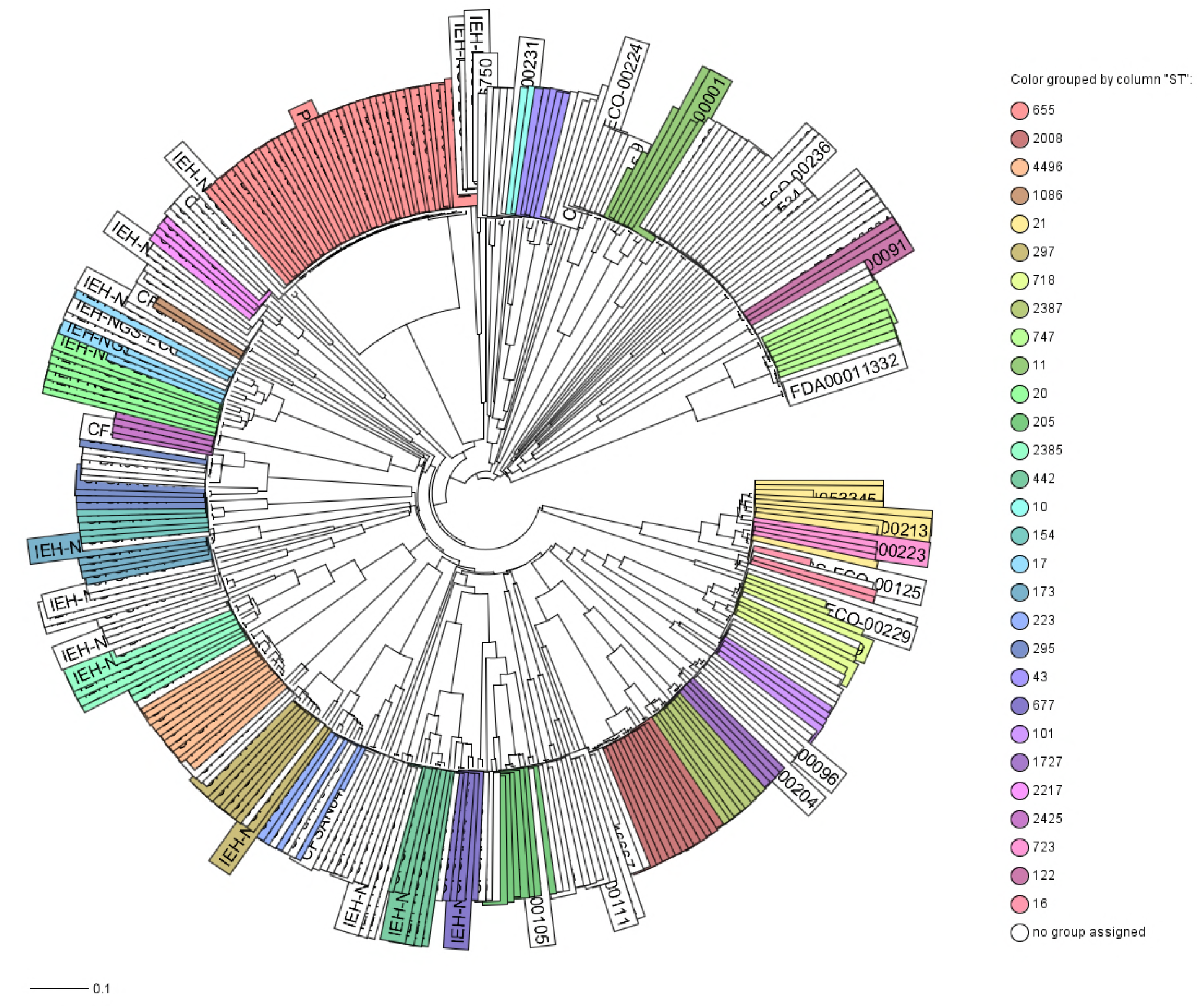
Phylogenetic relationships among the 276 STEC genomes of *E. coli* sequenced in this study by cgMLST analysis. Ridom SeqSphere+ (v5.0.0) identified 4,651core genes. The evolutionary history was inferred by using the Neighbor-joining (NJ) tree built using the genetic distance and showing the existence of many diverse clades with a complex evolutionary history. Strains are colored based on different STs as labeled.

The initial phylogenetic analysis [Neighbor-Joining (NJ) tree] based on gene differences (allele based) among these 276 STECs (Figure 1) revealed a complex evolutionary history with the existence of multiple, highly diverse genomic variants of strains isolated from RFFDA. Some of these genomes formed discrete groups and clustering was consistent with their ST (ex. all ST655 strains clustered together). A further analysis by a minimum spanning tree allows visualization of alleles differences between strains with the same ST that was not seen with the NJ tree (Figure 2).

**Figure 2.**
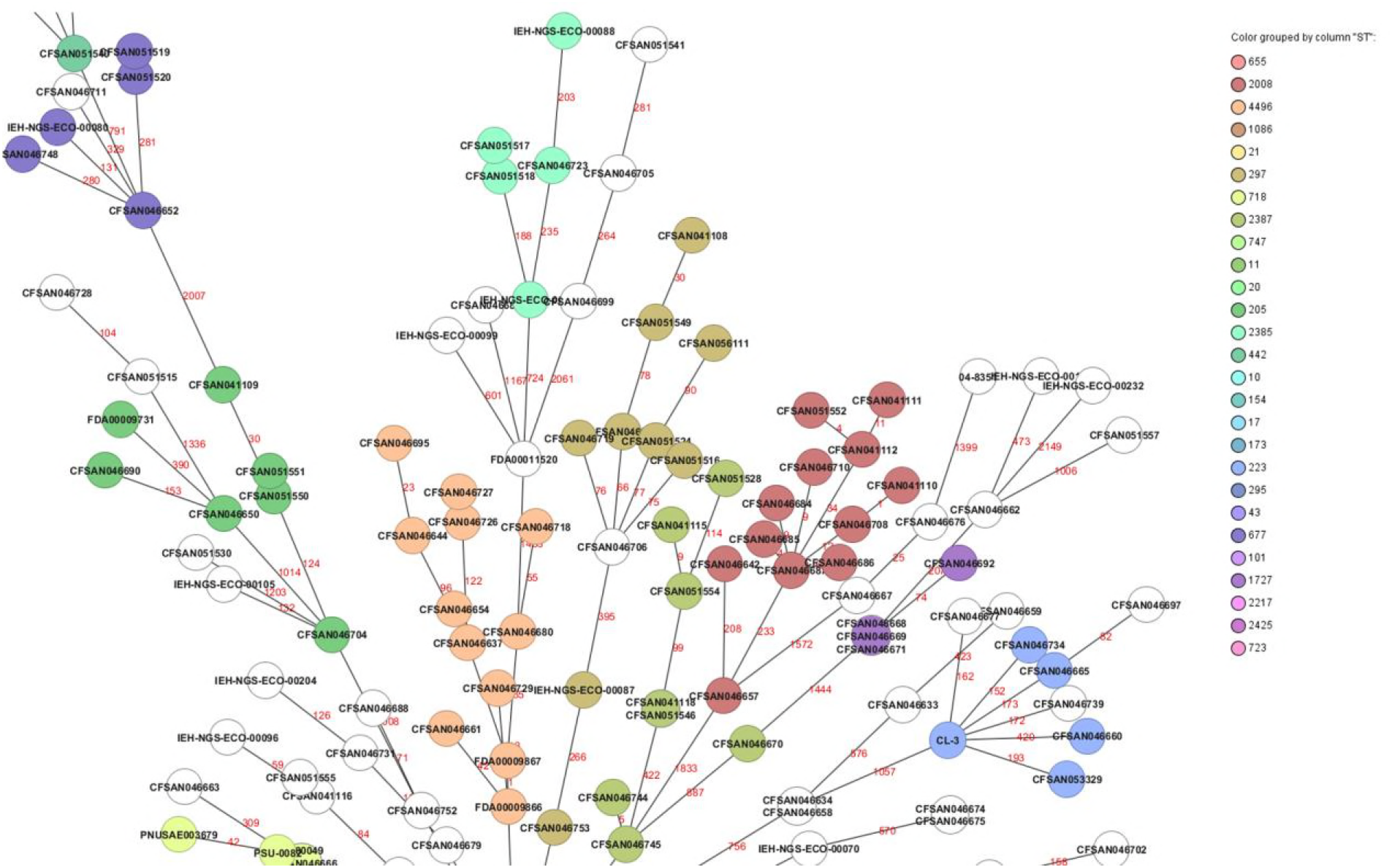
Snapshot of a minimum spanning tree showing the relationships among all different STECs. The numbers above the connected lines (not to scale) represent allele differences between strains belonging to the same ST. The isolates are colored based on different STs as labeled.

### *eae* positive Non-STEC strains virulence gene profiles

Among the 55 non-STECs (lacking either *stx* gene by *in silico* analysis) strains isolated from FDA regulated foods, we found 35 that were positive for the *eae* gene (Table 6). Most of them were classified as atypical EPEC (aEPEC) *eae*^+^ and *bfpA*^-^. Even though two of them (IEH-NGS-ECO-00094, and IEH-NGS-ECO-00100) carried *bfpA (eae^+^* and *bfpA^+^*) they were missing most of the common genes found in typical EPEC (Table 6, typical EPEC lineage 1 strain E2348/69). Therefore, we classified them as aEPEC. Virulence genes for ETEC, EIEC, and EAEC were not detected among the 331 sequenced *E. coli* strains (results not shown).

**Table 6.**
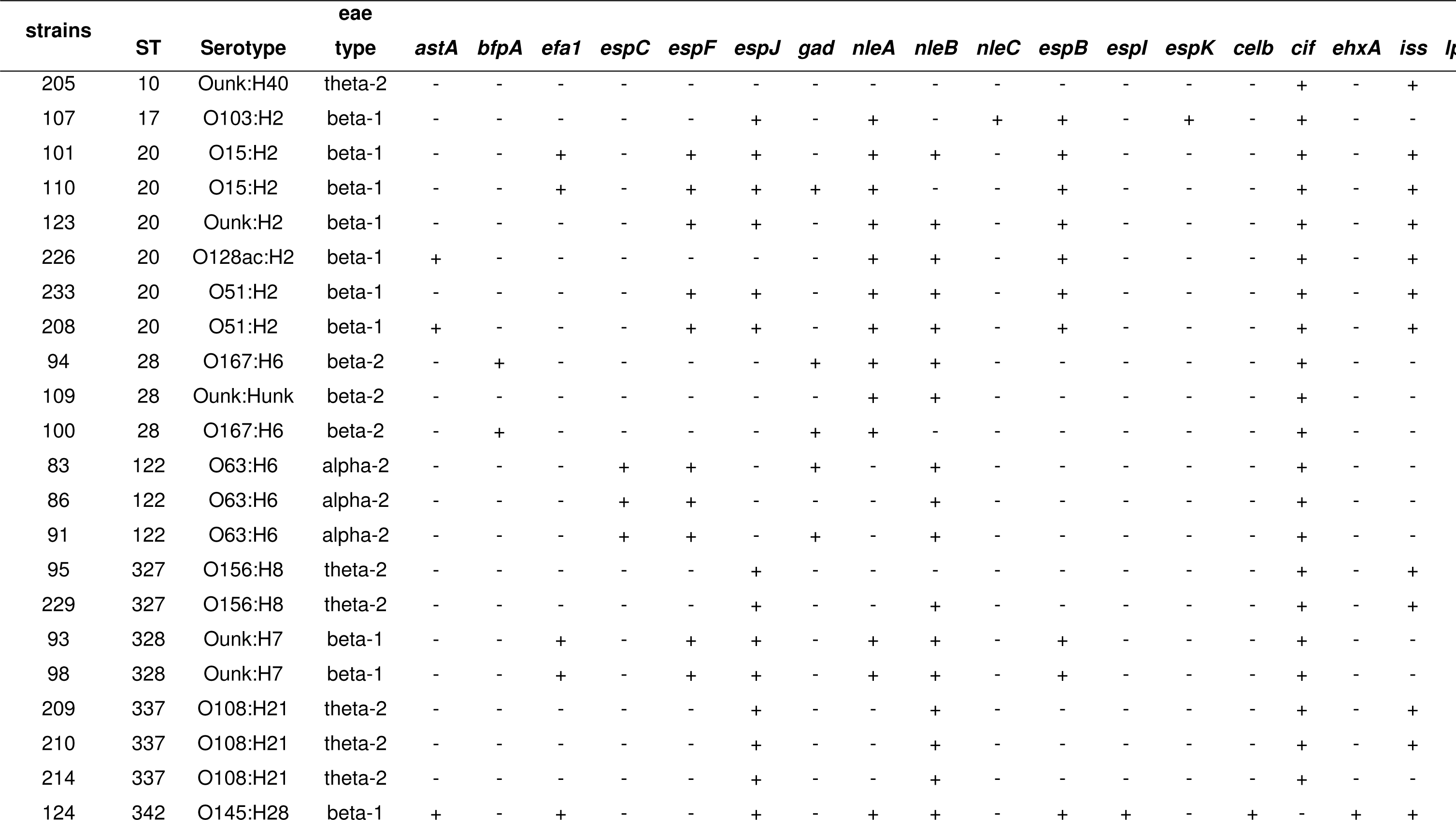
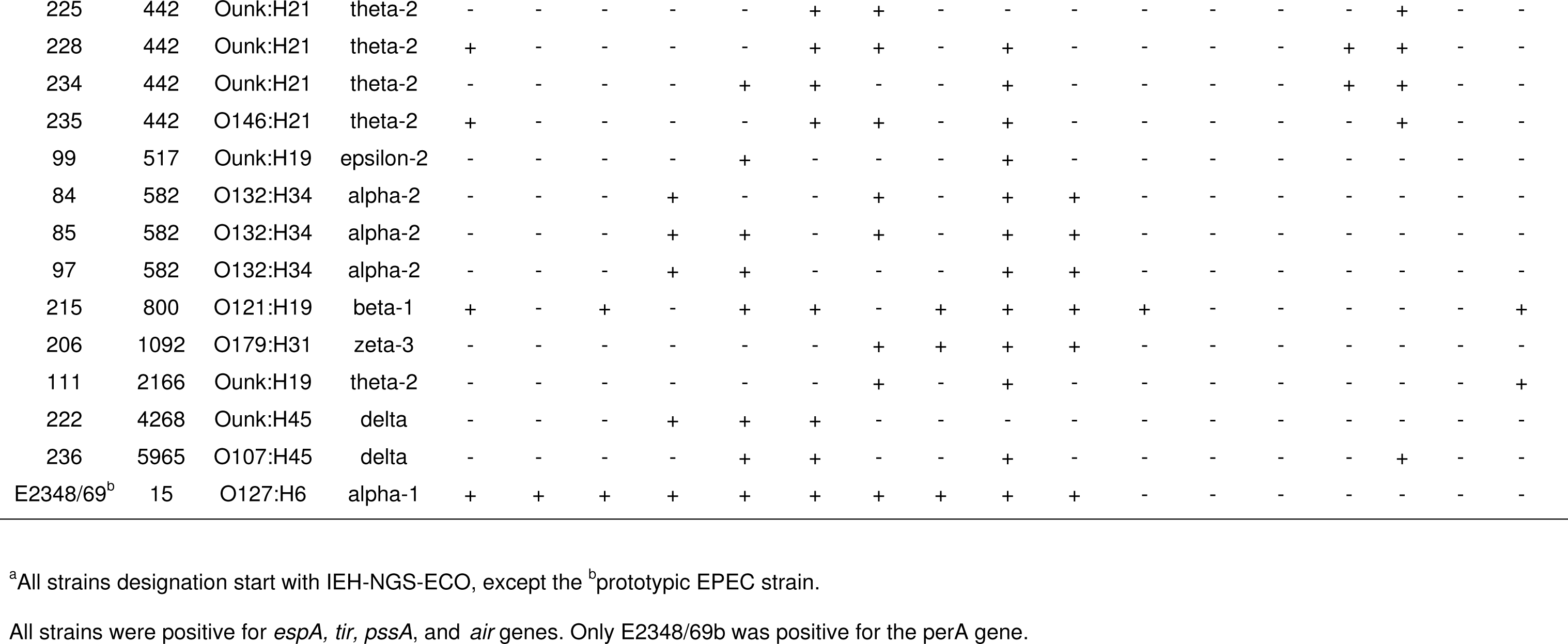
ST, serotype and virulence profile of intimin positive non-STEC isolated from FDA regulated foods (2003-2017).

## Discussion

STECs are the most dangerous among diarrheagenic *E. coli* for public health worldwide (5,7,23,43-45). Usually the most threatening STEC are those of O157:H7 serotype (46,47). However, in recent years there has been an increase in the occurrence of many non-O157 serotypes in humans associated with consumption of contaminated food, including produce and other FDA regulated products (2,48,49). Some studies have characterized STECs presence and their virulence potential from FDA regulated products (20,23,50). Most of the STEC isolated from those products have been only initially screened for the presence of some virulence genes using PCR (23,51). In the present study, we performed an in-depth analysis by whole genome sequencing of 331 presumptive STEC strains isolated from FDA regulated foods recovered during a period of 2003-2017 by two surveillance programs (FDA ORA, and MDP USDA) and other sources. STECs were isolated from 22 food commodities. It is worth mentioning that even though the sampling occurred in not all states, the food commodities had nationwide (or at least multistate) distribution.

The STEC analyzed in this study were isolated from a wide variety of foods (Table 1), with the majority isolated from spinach (32%), flour (21%), lettuce (13%), and cilantro (12%) samples during the period 2010-2017. The actual frequency of flour STECs should be assessed at a lower frequency of 9%; the spike observed in their frequency was due to the outbreak in flour in 2016, where most STECs (37 strains (66%) of total flour STECs) were isolated. A better reflection of the frequency of STEC isolated per food commodity, specifically produce, can be found in Feng and Reddy (2013) (23). Nevertheless, the presence of STECs in FRFDA per year remained relatively low, with a median of 30 isolates per year. As pointed previously, these variations in frequency of isolation can be due to seasonal variations, geographical variations, or even due to sporadic outbreaks as was observed for O121:H19 STEC strains isolated from flour in 2016 (23).

WGS revealed that these STECs were highly variable with the existence of 95 different sequence types (STs) and belonging to at least 81 different serotypes. Some serotypes could not be predicted and might be due to that the O type and H type genes were not present in the database used which includes the most frequent serotypes found in clinical cases. Most STs were observed only once while some others were observed more frequently. ST655 was observed up to 38 times among the STECs analyzed and it was because 37 of those STEC strains were recovered during the flour outbreak in 2016 (20). The majority of the STEC STs observed in this study [69/95-73%] had been reported as causing disease in humans, according to what was found in Enterobase (http://enterobase.warwick.ac.uk). Furthermore, of these potential human pathogenic STECs, strains belonging to 18 of those STs (19%) were additionally associated with strains causing EHEC-related illnesses (Table 3). Among the known ST associated with causing HC illnesses or HUS cases we found: ST21 and ST29 (O26:H11), ST11 (O157:H7), ST33 (O91:H14), ST17 (O103:H2), and ST16 (O111:H-), among others (44, 52-54).

Some samples have the same ST however they show differences in their virulence profile as well as their Shiga toxin gene content. For example, there were 5 strains that were ST10 and from these only 2 were classified as STECs, with one carrying *stx*1a while the other carried *stx*2a, both were negative for any of the attaching genes (*eae*, *saa,* and *subA* genes), therefore considered as low risk of causing infection in a healthy individual. This is an example that a single characteristic (e.g. ST or serotype) is not enough to make an inference of the potential pathogenic trait of any STECs (http://www.fao.org/documents/card/en/c/CA0032EN). The better way is to take all the information into consideration (ST, *stx* type, attaching genes, serotype, etc) in order to make a more informed prediction of the pathogenic potential of any STEC in conjunction with historical available data on clinical cases. For example, a strain of O113:H21 *stx*2a positive and that doesn’t possess *eae* but it has *subA, saa*, and *sab* genes, and has been linked to HUS cases (19), therefore, we can foresee that this strain might be harmful to humans. A similar analysis could be done in the case of any STEC that have all those attributes but that has not been linked to any human cases. Even though we cannot predict the actual outcome of an infection with this strain, it still warrants a warning about its presence in foods that are consumed raw as it is the case of fresh produce.

We tested for 95 known virulence genes (27) found in the most common *E. coli* pathotypes and did not find any genes present that would characterize the strains as STEC/EAEC//ETEC/EIEC hybrids. Among the adherence factors, *eae* and *subA* genes were found in 24% and 26% of the STEC strains, respectively. Strains that carry *eae* did not carry *subA* or *saa* genes, and viceversa, as perviously observed for STEC isolated from fresh produce (23). Regarding the presence of Shiga toxin type, there was great variation with most strains (67%) carrying only stx type 2, 19% carrying only stx type 1 while 15% carried both stx types. Among the stx type 2 there were 144 that were either a,d, or c, which are the stx2 types found among clinical cases (31,32,55-57) and that have specific trophism for humans (56). The remaining 40 STECs carrying *stx* type 2 alone were stx type (e, d/e and g) which have been described in animal reservoirs (58). The ones carrying both stx types were all stx1a. The remaining virulence genes were sporadically found with the most common *exhA* gene was found in 61% of the STECs, while *espP* was found in 43% of the STECs. These two genes can be found in the virulence plasmid and appears to participate in STECs infection in humans (9,11,13). In summary, 46 of the STECs analyzed in this study carried both *stx2*a and *eae* gene which is considered of elevated risk to human health (22,56).

We also confirmed the presence of antimicrobial resistance (AMR) genes using the DTU database with our modified protocol (Ridom) and find that their prevalence to be low at only 12%, although carrying multiple antimicrobial resistance genes. The presence of strains carrying multiple AMR genes is worrisome since they can be shared amongst other *E. coli* and could possible participate in the dissemination of AMR in their environments, as has been observed occurring for tetracycline genes in *E. coli* isolates from beef cattle (59), for colistin resistance (*mcr*-1 gene) through plasmid-mediated transfer (60), and for ampicillin resistance genes in *E. coli* in an infant treated with antibiotics (61).

Phylogenetic analysis by a custom cgMLST analysis of these 276 STECs confirmed the MLST *in silico* analysis, with many different defined clades among these STECs isolated from FRFDA. The cgMLST analysis is a fast method of analysis and provides an initial visualization of the relationships among the strains analyzed. Comparable results have been observed for establishing fast relationships among genomes from diverse bacterial pathogens (29,43,62-67). A further analysis using only the genomes of strains that are located within each individual or among selected clades can produce a more detailed evolutionary history, using single nucleotide analyses, which can help in determining to understand the potential source, phylogenetic nature, lineage, and timeline of transmission of each group, as has been shown for the ST36 lineage of *Vibrio parahaemolyticus* (68).

EPECs are the leading cause of infantile diarrhea in developing countries (70,71). Typical EPECs (tEPEC) have *eae* and *bfp* genes, and their main reservoir is humans (72). The *eae* gene is located in the chromosome, in the LEE operon, while the *bfp* operon is typically located in the large EPEC adherence factor (EAF) virulence plasmid (72). These tEPEC also carry the *perA* gene, which increases the expression of LEE elements (72,73). Interestingly, 17% of our presumptive STECs were shown by *in silico* analysis to be atypical EPECs (aEPEC). Among their unusual features are the absence of the EAF plasmid, and their reservoirs can be animals or humans (72). It is possible these aEPECs might had lost their phages upon culturing, as this pattern has been observed in clinical isolates of *E. coli* upon sub-cultivation (69). The aEPEC we observed in this study may have the capacity to produce A/E lesions, since they carried both the *eae* and *tir* gene, which are the effector and receptor necessary for the formation of the A/E lesion (74).

Our results suggest that finding aEPECs in food could be of particular concern, as these strains have the potential for acquiring the *stx* phage, as observed in the *E. coli* O104:H4 strain found in Germany (75). That strain was an entero-aggregative *E. coli* (EAEC) that had acquired an *stx*2a phage, and human illnesses that resulted during 2011 became the largest known HUS outbreak of STEC-related illness in the world (75). Similarly, anO26:H11 strain 21765, isolated in 2005 during a milk cheese outbreak in France (76) was shown to be an EPEC strain that had probably acquired a *stx*2a phage (27). In Gonzalez-Escalona et al (2016), the authors demonstrated that some strains of *E. coli* O26:H11 isolated from US cattle were phylogenetically more closely related to ST29 O26:H11 EHECs but t because these did not carry the *stx* phage, they would have been classified as EHEC-like by previous methods (77). Over the last five years, the analyses of thousands of *E. coli* genomes have revealed that so-called *E. coli* “hybrid strains” – strains that belong to one pathotype but acquire virulence markers, such as *stx* genes, from another pathotype – could be more common than previously believed. If this is the case, this suggests that environmental *E. coli* strains currently considered harmless could acquire the potential to pose risks to human health; for example, both aEPEC and STEC strains were isolated from foods such as flour, cilantro, lettuce, and kale (Table S1).

We are heading to a new phase in surveillance of STECs in the US by using a genomic monitoring approach and our STEC sequences from FRFDA provides a solid foundation to build upon (78)( https://www.cdc.gov/pulsenet/pathogens/wgs.html). There already exists a database that achieve the first goal of source tracking by using core genome information (NCBI pathogen detection tool), but there is a need for improved databases that allow for fast analysis of the WGS data for detecting virulence genes, phages and plasmids content, as well as antimicrobial resistance genes.

In conclusion, STECs were isolated from diverse FRFDA food sources during the period study. The contamination frequency was relatively low (average 30 STEC strains isolated per year). However, fifty percent of the STECs analyzed in this study carried either a combination of *eae* plus *stx*, or *subA* plus *stx*, therefore being potentially pathogenic to humans. Moreover, those STECs carried most of the virulence genes described for STECs causing infections with a diverse range from HC (e.g. ST655 O111:H19 strains) to HUS (e.g. ST21 O26:H11 strains) (20,44). Some others have not been described as causing disease in humans but have the potential to do so (e.g. ST342 O5:H-unknown strains) since they carried all virulence genes described in pathogenic strains (*stx1a*, *eae*-beta1, *exhA*, *tir*, and many of the T3SS effectors and non-LEE effectors) (Table 4). Nonetheless, the determination of the presence of STECs in FRFDA with potential to cause disease in humans reinforce the need to continue surveillance for this important pathogen which is of importance for food safety and public health. Furthermore, the availability of these genomes could provide early warnings of food contamination from cattle or other animals, since some of the STEC isolated were carrying stx2e that have been usually observed causing edema in pigs (79) and are considered as probably non-pathogenic to humans (56). Here we showed that WGS enabled comparisons across isolates to establish phylogeny, help in identification of antibiotic resistance by monitoring the presence of antimicrobial resistance genes, and determined the presence of known virulence genes that have been linked with illnesses. A freely accessible database of high-quality reference genome sequences of FRFDA was previously unavailable. Future food safety investigations will benefit from the comparisons made possible by WGS databases like ours as it allows for the monitoring of the recurrence and emergence of strains in the food supply. It is our goal to help develop databases that will allow for fast source tracking and accurate categorization (low risk or high risk) of STECs food isolates in a more comprehensive manner.

## ACKNOWLEDGMENTS

This work was supported by the FDA Foods Science and Research Intramural Program. We also want to thank Sabina Lindley for assistance in sequencing some of these *E. coli* strains. The authors thank Lili Fox Vélez for her helpful comments on this manuscript.

---

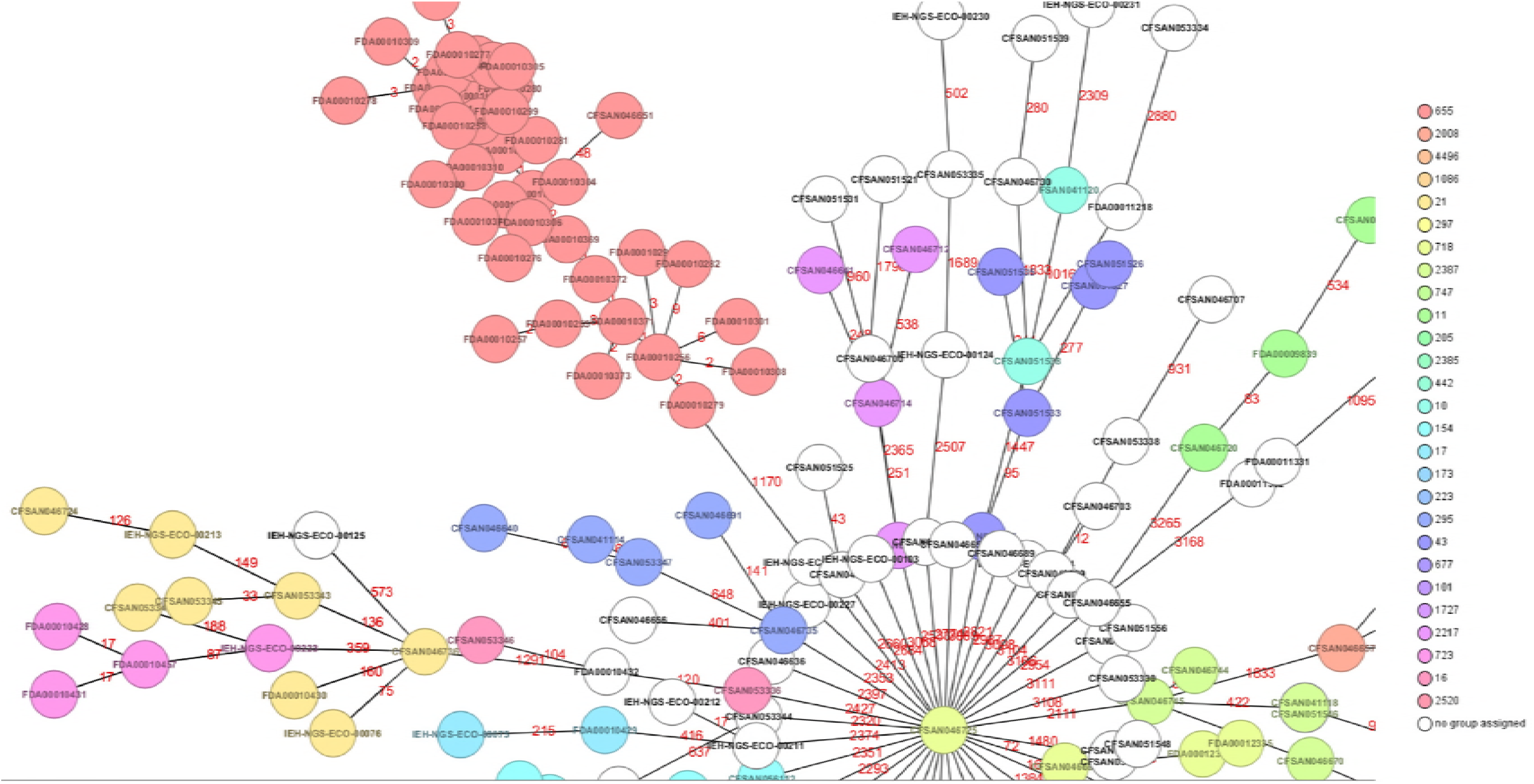

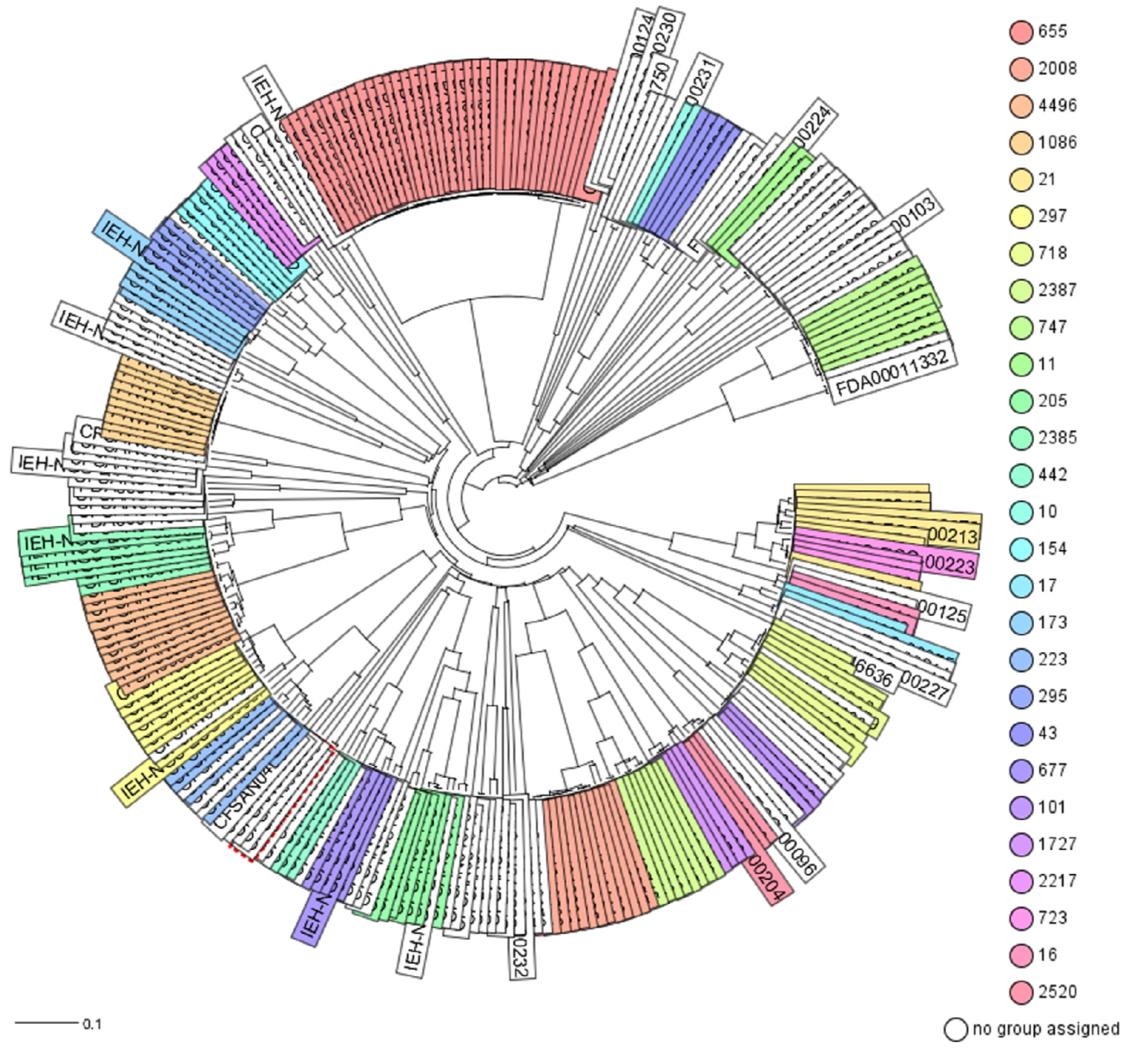

